# Training qualitatively shifts the neural mechanisms that support attentional selection

**DOI:** 10.1101/091413

**Authors:** Sirawaj Itthipuripat, Kexin Cha, Anna Byers, John T. Serences

**Affiliations:** Neurosciences Graduate Program University of California, San Diego, La Jolla, California 92093; Department of Psychology University of California, San Diego, La Jolla, California 92093; Kavli Institute for Brain and Mind University of California, San Diego, La Jolla, California 92093

**Keywords:** Attention, Training, Gain, Noise, Contrast discrimination, Signal detection Theory, EEG

## Abstract

Attention supports the selection of relevant sensory information from competing irrelevant sensory information. This selective processing is thought to be supported via the attentional gain amplification of sensory responses evoked by attended compared to unattended stimuli. However, recent studies in highly trained subjects suggest that attentional gain plays a relatively modest role and that other types of neural modulations – such as a reduction in neural noise – better explain attention-related changes in behavior. We hypothesized that the amount of training may alter neural mechanisms that support attentional selection in visual cortex. To test this hypothesis, we investigated the influence of training on attentional modulations of stimulus-evoked visual responses by recording electroencephalography (EEG) from humans performing a selective visuospatial attention task over the course of one month. Early in training, visuospatial attention induced a robust attentional gain amplification of sensory-evoked responses in contralateral visual cortex that emerged within ~100ms after stimulus onset, and a quantitative model based on signal detection theory (SDT) successfully linked this attentional gain amplification to attention-related improvements in behavior. However, after training, this attentional gain amplification of visual responses was almost completely eliminated and modeling suggested that noise reduction was required to link the amplitude of visual responses with attentional modulations of behavior. These findings suggest that the neural mechanisms supporting selective attention can change as a function of training and expertise, and help to bridge different results from studies carried out in different model systems that require substantially different amount of training.

## Introduction

Selective attention mediates the processing of sensory information so that relevant information is preferentially processed over irrelevant information. Over the last several decades, multiple electrophysiological and neuroimaging studies in humans and non-human primates (NHPs) have shown that attention selectively increases the amplitude of visual responses evoked by attended stimuli compared to those evoked by unattended stimuli (i.e., attentional gain) (1–31). Particularly in studies that use human participants, the magnitude of these attentional gain changes has been shown to be tightly associated with attentional modulations of behaviorally measured perceptual sensitivity (7,11,14). For example, human electroencephalography (EEG) studies demonstrated that the attentional gain amplification of neural responses in contralateral visual cortex that began within ~100ms post-stimulus were highly correlated with subjective reports of perceived contrast of visual stimuli (11) and with improvement in target detection performance (7). Moreover, it has recently been shown that attention-induced changes in psychophysically-measured perceptual contrast discrimination performance could be accurately predicted by the observed amount of attentional gain amplification of these visual responses (14). Taken together, these EEG findings suggest that the attentional gain amplification of neural responses in visual cortex has a significant impact on perception and behavior during behavioral tasks that require selective attention.

Despite this support for the importance of attentional gain, a growing number of recent studies suggest that other types of neural modulations in visual cortex more closely track perceptual performance. For example, electrophysiological studies in NHPs have demonstrated that, in addition to enhancing gain, attention can also reduce trial-by-trial variability in single neuron spike rates as well as pairwise correlations between neurons (32–40). Moreover, these modulations of neuronal noise may improve the signal-to-noise ratio of sensory codes more than attentional gain (33,38) and are also more closely correlated with changes in behavioral performance (33). While these two attention mechanisms are not mutually exclusive, the relative contribution of each type of modulation to behavioral performance is hard to evaluate given large differences in methodologies and animal model systems employed across different studies (18,41–44). One particularly salient difference concerns the amount of training that subjects receive on behavioral tasks before neural data is collected. For example, studies that tend to reveal attentional gain as a predominant attention mechanism typically used human participants trained for brief periods of time (typically less than one hour) (8–11,13). On the other hand, studies that support the importance of other mechanisms such as noise reduction typically used monkeys trained for many months (32,33,38). Thus, understanding the impact of training on the mechanisms that support selective attention is important for generalizing results across tasks and model systems that involve different levels of training and expertise.

To directly test the influence of training on the mechanisms underlying attentional selection, we had human participants perform a selective visuospatial attention task for over one month. Throughout training we concurrently measured their psychophysical performance and brain activity using electroencephalography (EEG), which is an advantageous approach for several reasons. First, human participants require less practice with complex tasks compared to other model systems, which enabled us to immediately acquire measures of neural activity with relatively little initial training and to continue tracking neural activity over the course of one month. Second, several past studies have used a positive-going event-related potential (ERP) that peaks about 100ms after stimulus onset (termed the P1 component) to measure the gain of population-level responses in extrastraite visual cortex (6,8–14,31,45–47). Importantly, the P1 component has been shown to be sensitive both to stimulus intensity (e.g., contrast and luminance) and attentional modulations (14,46). Accordingly, we focused on examining the impact of training on the attentional gain of the P1 component in human participants performing a contrast-discrimination visuospatial attention task. We expected to observe a robust attentional gain amplification of the P1 component early in training based on previous results (6–14,47). However, if training leads to a transition from attentional gain to another mechanism such as noise reduction, then attentional modulations of the P1 component should be reduced over time (Fig. 1). To link attentional modulations of the P1 with behavior at different points in training, we also adopted a standard quantitative model, based on signal detection theory (SDT), to infer whether attentional gain or noise reduction best predicted behavior (14,25,26,48–50). The results are consistent with a qualitative shift from attentional gain to noise reduction over time, suggesting that extended training can alter the mechanisms that support selective information processing in visual cortex.

**Fig. 1.**
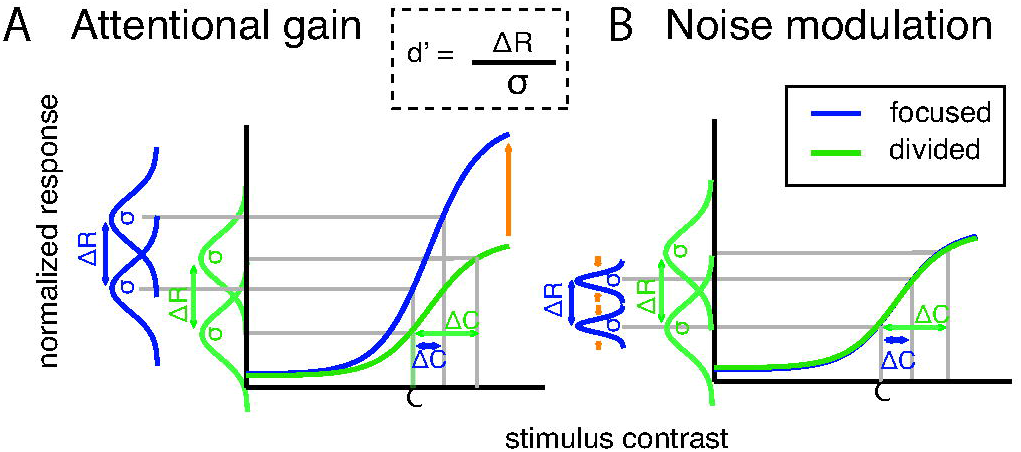
Competing attention mechanisms. (A) Attentional gain mechanisms posit that selective attention enhances early visual responses, which increases the signal-to-noise (SNR) ratio and perceptual sensitivity (d’). (B) Noise modulation models hold that attention impacts SNR via changes in neuronal variability. According to the signal detection theory (SDT), perceptual sensitivity (d’) increases via increasing gain (ΔR) and via reducing noise (σ). We hypothesized that training might qualitatively alter the neural mechanisms that support attentional selection. Specifically, we predicted a qualitative shift from attentional gain (A) to noise reduction (B) over time.

## Results

### Behavioral Results

Human participants performed a two-interval-forced-choice (2IFC) contrast discrimination task (Fig 2A). We used this task to make contact with previous studies in both humans and NHPs that have employed similar paradigms (14,25,26,32,35). On each trial, subjects were cued to attend to either the left or right lower visual quadrant (termed focused attention), or to attend to both locations (termed divided attention). The cue was followed by two successive stimulus intervals, and each interval contained one sinusoidal Gabor stimulus to the left and one to the right of fixation. In the focused attention condition, the two successive stimuli at the cued location were always rendered at different contrast values. In one of the two stimulus presentation intervals, the contrast value of each stimulus was pseudo-randomly drawn from 0%-61.66% Michelson contrast. We refer these contrast values as ‘pedestal’ contrast values. For the other stimulus interval, we added a slight contrast increment to the pedestal contrast value of one of the two stimuli and participants then had to report whether the first or the second interval contained the stimulus with a higher contrast value. At the uncued location, the two successive stimuli were always rendered at the same contrast value drawn from 0%-61.66% Michelson contrast. We refer to the stimuli presented in the cued location as the ‘focused target’ stimuli and the stimuli presented in the un-cued location as the ‘focused non-target’ stimuli. In the divided attention condition, both locations were equally likely to contain the contrast change, yielding the ‘divided target’ and ‘divided non-target’ stimuli, which contained and did not contain a contrast change, respectively. The main dependent measure was the change in contrast (Δ*c*) from each pedestal contrast that was required to achieve an accuracy level of 76% (a d’ of approximately 1). Importantly, this method allowed us to derive both neurometric and psychometric response functions so that we could directly link modulations in neural responses with modulations in behavior (see also:14,25,26,48).

**Fig 2.**
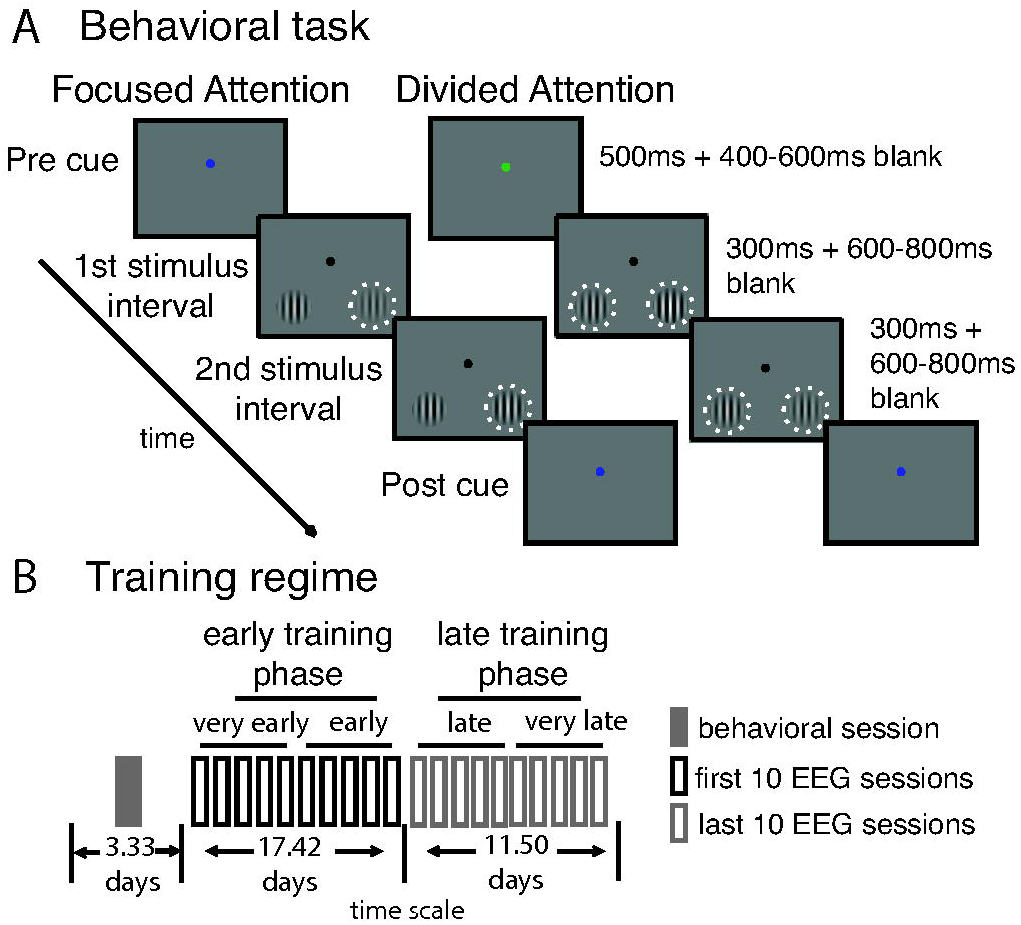
Task design. (A) A two-interval force choice (2IFC) contrast discrimination task that required either focused or divided visuospatial attention. (B) Training regimen: 12 participants completed one session of behavioral training and then completed 20 sessions of simultaneous behavioral testing and EEG recording over an average of 32.25 days.

To evaluate the effects of training on attentional modulations, 12 human participants participated in 20 EEG sessions (1-2 sessions each day) over the course of approximately one month (Fig 2B). In each experimental session, we estimated the incremental contrast value (Δ*c*) required to reach criterion performance at each pedestal contrast and attention condition (Fig 3A; mean hit rate: 76.6% ±SEM 0.3%, yielding d’ = ~1). From this point on, in the main text, we discuss the data divided into an early training phase (first 10 sessions) and a late training phase (last 10 sessions). However, note that similar results were observed with the data divided into 4 phases (5 EEG sessions in each phase) or 10 phases (2 EEG sessions each phase; see S1 supporting Information).

**Fig 3.**
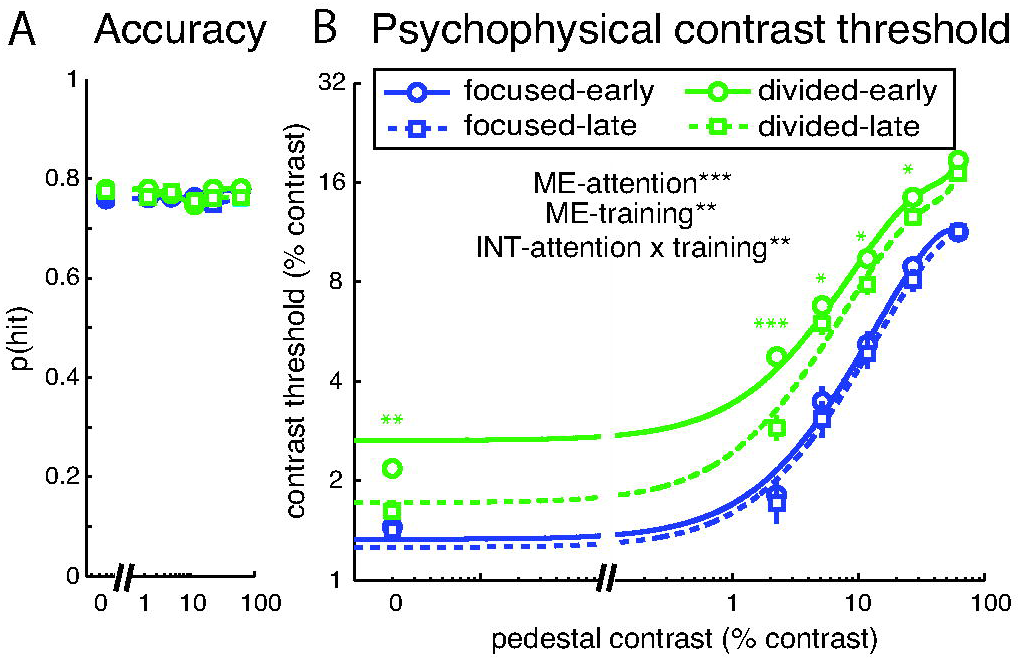
Behavioral results. (A) The hit rate was fixed at ~76% across all conditions so that contrast discrimination thresholds could be measured as a function of attention and training. (F) Contrast discrimination thresholds (Δc) were lower on focused attention trials compared to divided attention trials. Training also led to lower discrimination thresholds, specifically when attention was divided and the pedestal stimuli had low to medium contrast levels. Error bars represent within-subject standard error of the mean (SEM). Black ** and *** represent significant main effects (ME) and interactions (INT) with p< 0.01 and p < 0.001. Green *, **, and *** represent pairwise differences between contrast thresholds in the divided attention condition across training phases with p <0.05, p<0.01, and p<0.001, respectively. Also see data divided into 4 and 10 training phases in S1 Fig 1.

Fig 3B illustrates the psychophysical contrast thresholds (Δc) required to achieve a fixed hit rate as a function of pedestal contrast in the focused and divided attention conditions (to produce a threshold versus contrast curve, or TvC). In line with previous studies, Δc increased as a function of contrast (F(5, 55) = 143.38, p <0.001; 14,26,52–55). In addition, Δc was smaller in the focused attention condition compared to the divided attention condition at all contrast levels (F(1, 11) = 137.95, p<0.001; all t(11)’s ≥5.03, all p’s < 0.001, Holm-Bonferroni correction, one-tailed; 14,26,53). Similar results were observed with the data divided into 4 phases or 10 phases (S1 Fig 1). Training also decreased Δc (F(1, 11) = 9.85, p = 0.009). However, the training effect on Δc was driven primarily by improved performance in the divided attention condition (F(1, 11) = 11.96, p = 0.005), particularly when the pedestal contrast levels were low (0% contrast: t(11) = 2.86, p = 0.008; 2.24% contrast: t(11) = 4.70, p < 0.001, Holm-Bonferroni correction, one-tailed). The relatively small effect of training in the focused attention condition is consistent with previous studies that used similar contrast discrimination tasks (55,56).

### EEG Results

The early visual system has a contralateral mapping between external stimuli and their cortical representation, such that stimuli presented in the left visual field evoke responses in right occipital cortex and vise versa. However, EEG has relatively coarse spatial resolution, thus ERPs recorded over the occipital lobe typically reflect a mixture of responses evoked by both stimuli (unless a 0% contrast stimulus was presented). To better isolate the stimulus-evoked responses associated with stimuli presented on the left and right sides of space, we first subtracted the averaged ERPs on trials that had a 0% contrast stimulus in the visual field contralateral to a given electrode from the averaged ERPs on trials where visual stimuli were rendered at each of the other contrast levels (Fig 4; see similar methods in 11,14,58). This subtraction method was performed separately for each stimulus contrast level, each attention condition and each training phase. This method helped not only to isolate evoked responses associated with a single stimulus from a bilateral stimulus array, but also to control for any spatially non-specific anticipatory effects associated with the presentation of attention cues (11,14,42,57). As a result, we observed clearly lateralized visual P1 components that scaled with increasing stimulus contrast. As shown in Fig 5, the P1 components peaked in the contralateral posterior-occipital electrodes ~80-130ms post-stimulus, consistent with previous ERP studies (6,8–14,31,45–47).

**Fig 4.**
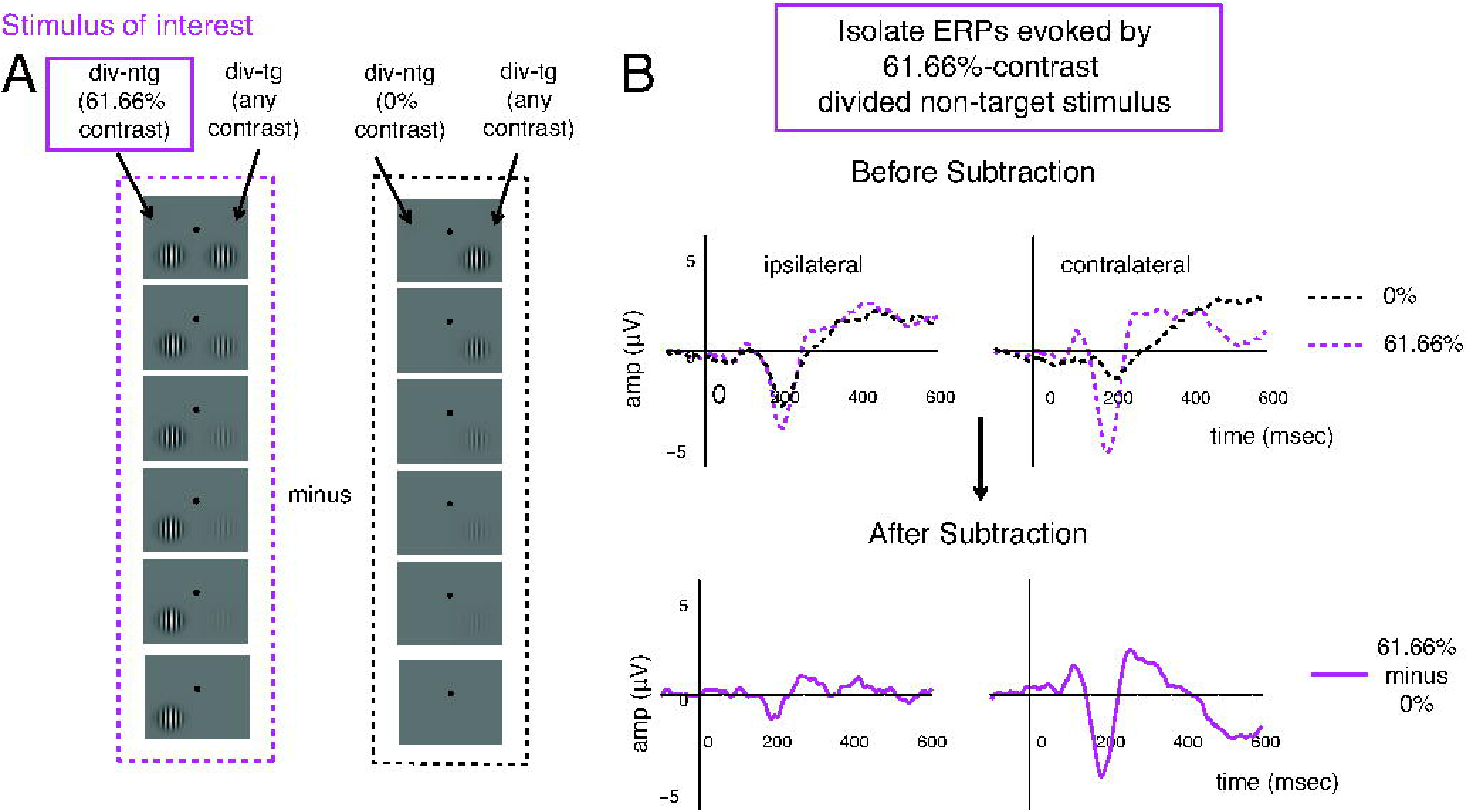
An example of the ERP subtraction method. (A) Left column (in purple): schematic of a 61.66% contrast divided non-target stimulus presented in the left hemifield (termed the stimulus of interest) and paired with target stimuli rendered in all different contrasts (down the rows). Right column (in black): the divided non-target 0% contrast stimulus in the left hemifield, paried with the same set of target stimuli in the right hemifield. (B) In this case, the ERP response evoked by the left divided non-target stimulus of 0% contrast (A, right; B, top, black dotted traces) was subtracted from the ERP response evoked by the left divided non-target-stimulus of 61.66% contrast (A, left; B, top, dotted purple traces), resulting in the baseline-subtracted ERP response (B, bottom, solid purple traces). A similar subtraction was done to compute the ERPs associated with stimuli of interest rendered at all other contrasts. Note that the stimulus paired with the stimulus of interest (in this case, the right divided target stimulus) could have any of six contrast values. Therefore, this method amounts to subtracting out the mean response to all ipsilateral stimuli.

**Fig 5.**
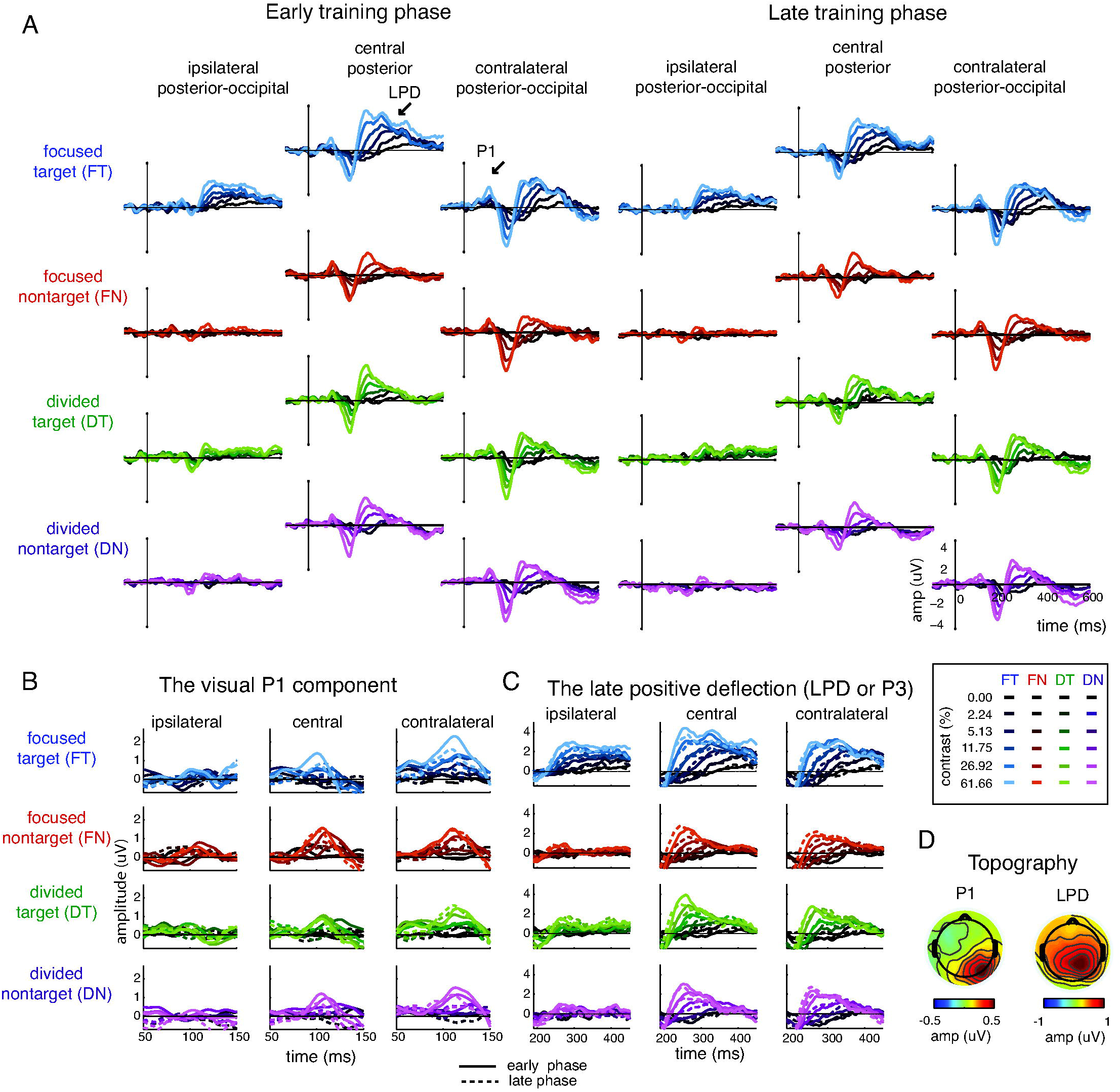
ERP results. (A) Extracted ERP traces evoked by focused targets, focused non-targets, divided targets, and divided non-targets across early and late training phases. The shading of the colors represents the contrast level of the stimulus (dark to bright colors represent low to high contrast levels). The ERP subtracting method, which helped isolating the ERPs evoked by the stimulus of interested from the bilateral stimulus array and helps minimizing cued-related anticipatory responses, is illustrated in Fig 4. (B) The zoom-in figure of the visual P1 component. (C) The zoom-in figure of the late positive deflection (LPD or P3). (D) Topographical maps of the P1 and the LPD component collapsed across all experimental conditions. The left and the sides of the topographical map depict the response in electrodes that are ipsilateral and contralateral to the stimulus of interest, respectively.

### Attentional gain amplification of the visual P1 component was attenuated with training

Fig 6A shows the P1 mean amplitude from 80-130ms post-stimulus over the contralateral posterior-occipital electrodes as a function of stimulus contrast, yielding contrast response functions (CRFs) for focused target/non-target and divided target/non-target conditions (see the logic for selecting analytic electrodes and time window in Materials and Methods). We then characterized the shape of the CRF in each condition using a Naka-Rushton equation (see Materials and Methods, Equation 1) to estimate the maximum response and the horizontal position of the CRF along the x-axis. Consistent with a recent report (14), we observed an increase in response gain or the maximum response of the P1-based CRFs with focused attention early in training (Fig 6A left). However, later in training, the attentional gain amplification of the P1-based CRFs was abolished (Fig 3A right). This led to a significant three-way interaction between training (early/late), attention (focused/divided), and stimulus type (target/non-target) on the maximum response (Fig 6B top; p = 0.036, resampling test, two-tailed; see Materials and Methods). This significant interaction was driven by the presence of a significant main effect of attention and a significant interaction between attention and stimulus type on the maximum response during the early training phase (p = 0.003 and p = 0.009, respectively, resampling test, two-tailed), but no main effect of attention or stimulus type and no interaction between these factors during the late training phase (p = 0.89, p = 0.54, and p = 0.83, respectively). This lack of attentional gain amplification after training was due to a selective reduction in the maximum response in the focused target condition (p = 0.013, two-tailed), accompanied by no changes in either the focused non-target, divided target, or divided non-target condition (all p’s ≥ 0.51). Follow-up analyses also revealed that the reduction in the attentional gain of the P1 occurred gradually in the focused target condition across training phases when the data are divided into 4 and 10 training phases (S1 Fig 2). The fact that the training-related change in P1 amplitude was specific to the focused target condition suggests that training specifically impacted neural modulations related to the deployment of focused attention. Moreover, the specificity of these modulations indicates that training-related changes in P1 amplitude were not due to general low-level sensory/perceptual learning effects since training did not impact the magnitude of the P1 associated with any other condition (i.e. divided target and non-targets and focused non-targets). While training had a significant impact on the attentional modulation of the maximum response, the contrast at which the response reached half maximum, which governed the horizontal positon of the CRFs on the x-axis, was unchanged (Fig 6B bottom). There were no main effects of training (p = 0.95), attention (p = 0.91), or stimulus type (p = 0.78), and no significant interactions between any of these factors (all p’s ≥ 0.13, resampling tests). Collectively, these results suggest that training primarily impacts the degree to which attention amplifies the maximum response of early stimulus-evoked responses.

**Fig 6.**
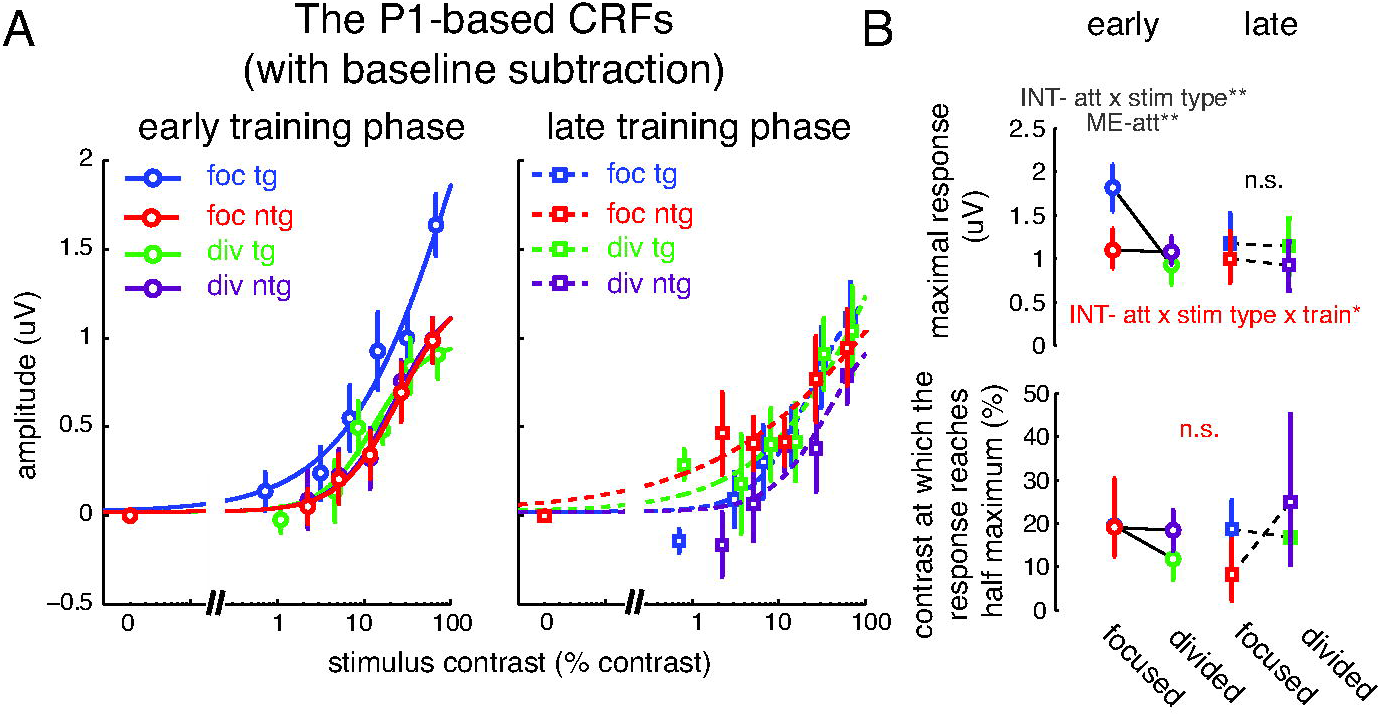
The P1 component with baseline subtraction. (A) The contrast response functions (CRFs) based on the amplitude of the P1 component averaged over the contralateral posterior-occipital electrodes from 80-130ms post-stimulus. During the early training phase, there was a robust attentional gain amplification of the P1 component on focused attention trials compared to divided attention trials (left panel). However, no attention-related gain modulations were present during the late training phase (right panel). (B) Corresponding maximal response and contrast at which response reaches half maximum for the P1-based CRFs. Error bars in (A) represent within-subject SEM. Error bars in (B) represent the 68% CIs. * and ** represent significant main effects (ME) and interactions (INT) with p <0.05 and p<0.01, respectively. Red signs show significance when data were compared across training phases. Gray signs show significance when data were compared within each training phase. See data divided into 4 and 10 training phases in S1 Fig 2 and data without baseline subtraction in Fig 7. Note that the contrast values on the x-axis are not exactly the same across target and non-target conditions because, in the target conditions, we used the averaged contrast values between the pedestal and incremental stimuli.

For comparison, we also analyzed the P1-based CRFs without subtracting the baseline activity. As illustrated in Fig 7A, the results were qualitatively similar to the results obtained using the subtraction method (Fig 6A). First, we fit the Naka-Rushton equation (Equation 1) to characterize the CRFs, but this time we included an additional free parameter to account for baseline differences between conditions. We then performed a nested model comparison to assess the goodness of fit between the model that allowed baseline parameters to change freely and the model that fixed the baseline parameter across all experimental conditions (see Material and Methods). This analysis revealed that allowing baseline parameters to change freely did not significantly improve the goodness of fit (F(7, 15) = 0.47, p = 0.99, nested-test). This suggests that baseline parameters associated with the P1-based CRFs did not change with attention or with training. Moreover, we observed the same pattern of response modulation with attention during the early training phase that dissipated after extended training (Fig 7B top): there was a significant main effect of attention and a significant interaction between attention and stimulus type on the maximum response during the early training phase (p < 0.001 and p = 0.033, respectively, resampling test), but no main effect of attention or stimulus type and no interaction between these factors during the late training phase (p = 0.91, p = 0.15, and p = 0.27, respectively). In addition, the horizontal shift along the x-axis was unchanged across experimental conditions and training sessions (all p’s ≥ 0.44; Fig 7B bottom).

**Fig 7.**
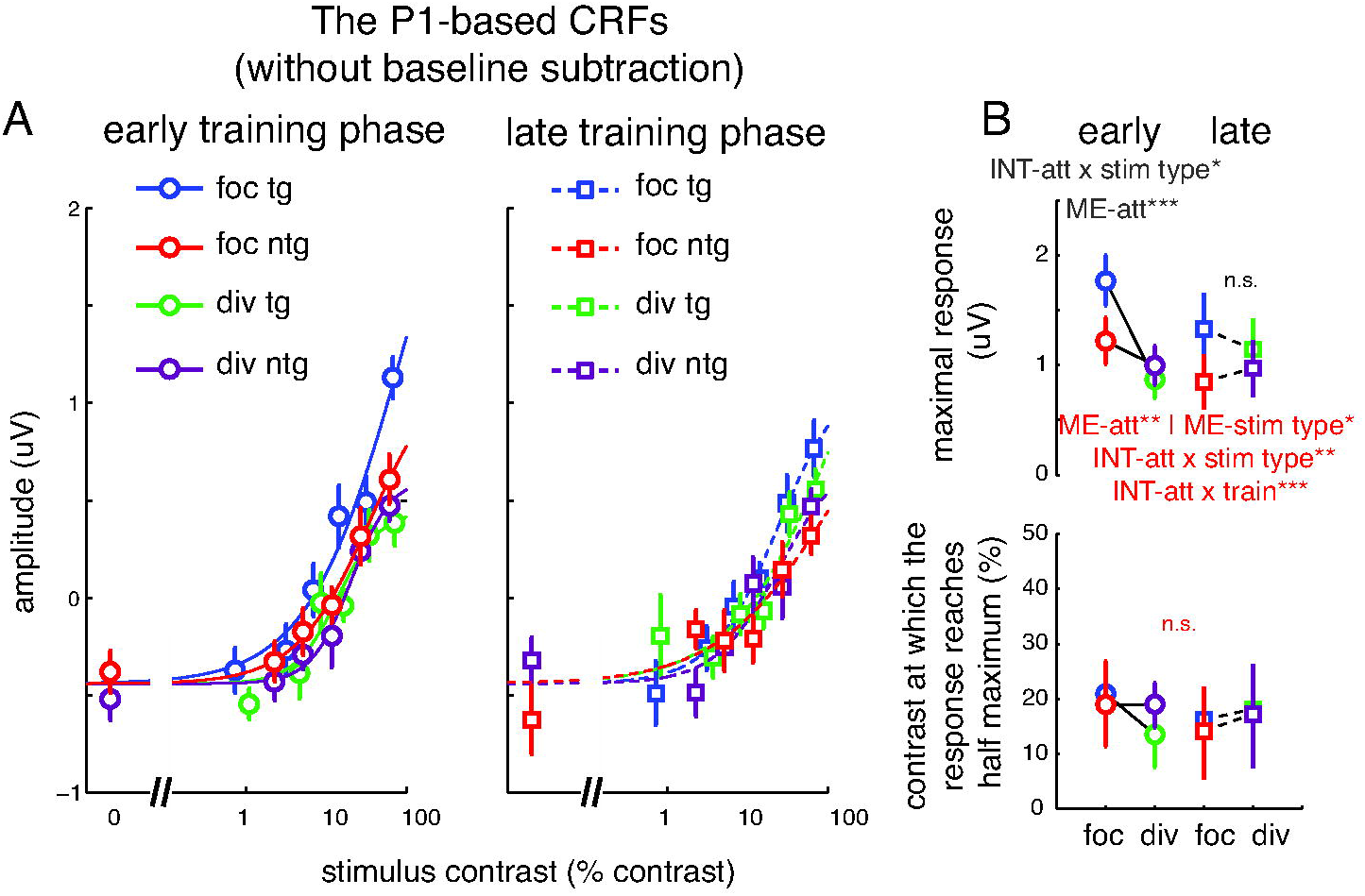
The P1 component without baseline subtraction. (A) The P1 CRFs based on ERPs without the baseline subtraction. (B) Corresponding maximal response and contrast at which response reached half maximum for the P1-based CRFs. Overall the results are consistent with the P1 results with baseline subtraction (Fig 6), where a gain modulation of the maximal response was observed during the early training phase, but this attentional gain modulation disappeared after extended training. The main difference between this result and the baseline-subtracted result is that the baselines of the P1 CRFs across all experimental conditions are shifted down and are negative. This is due to an early negative component induced by the stimulus that was ipsilateral to the electrode of interest. Error bars in (A) represent within-subject S.E.M. Error bars in (B) represent the 68% CIs. *, **, and *** represent significant main effects (ME) and interactions (INT) with p <0.05, p<0.01 and p <0.001, respectively. Red signs show significance when data were compared across training phases. Black signs show significance when data were compared within each training phase. Note that the contrast values on the x-axis are not exactly the same across target and non-target conditions because, in the target conditions, we used the averaged contrast values between the pedestal and incremental stimuli.

### Quantitative modeling based on signal detection theory (SDT) suggests a transition from gain to noise reduction after training

Overall, the P1 results suggest that while attentional gain is a prominent mechanism that supports attentional selection early in training, the absence of attentional gain amplification later in training suggests that training may alter the neural mechanisms that support attentional selection. Recent empirical and modeling evidence has been reported to supported alternative mechanisms including noise modulation (32– 36,39,40,58) and efficient read-out mechanisms (25,26). To evaluate these alternative accounts, we adopted a quantitative modeling framework based on SDT that can evaluate the impact of gain and noise modulations on behavioral performance (14,25,42,43,48). Later, we also evaluate the contributions of the efficient read-out mechanism in relation to gain and noise modulation mechanisms (25,26).

As illustrated in Fig 1 and Equation 3, The SDT-based model posits that the perceptual sensitivity (or d’) is determined by the difference between the mean responses (ΔR) evoked by two different stimuli (e.g., the pedestal and the increment stimuli) divided by the trial-by-trial variability of those responses (σ) (14,25,26,43,48–50). If attention operates solely via an attentional gain mechanism, it will increase ΔR, and hence increase d’ (Fig 1 left). Alternatively, if attention operates solely via a noise modulation mechanism, a reduction in σ is expected so that the overlap between the two response distributions becomes smaller and discrimination becomes more accurate (Fig 1 right). Thus, this modeling framework can be used to indirectly infer the relative contribution of attentional gain and noise modulations in situations where direct measures for neuronal noises are not available (14,25,26). According to this modeling framework, when d’ is fixed and there is a decrease in psychophysical contrast thresholds (Δc) with attention, a model based solely on changes in attentional gain (termed gain model) would predict an increase in the maximum response of the neural CRFs in the focused compared to divided target conditions. In turn, if these changes in gain are not sufficient to explain behavior, then the model will incorporate changes in σ (termed noise model) to improve the link between neural CRFs and behavior.

Consistent with a recent study (14), the gain model effectively linked changes in contrast thresholds and changes in the slope of the P1-based CRFs during the early training phase (compare black curve and blue circles in the middle panel of Fig 8A, R^2^ = 0.942). Moreover, the noise model did not significantly improve the fit between the behavioral data and the P1-based CRFs compared to the gain model (compare red and black lines in Fig 8A; R^2^ = 0.943, F(1, 8) = 0.12, p = 0.74, nested test). This suggests that the attentional gain mechanism can sufficiently account for the relationship between attentional modulations in neural and behavioral data early in training. Later in training there was no attentional modulation of the P1-based CRFs but there was still an improvement in behavior (Fig 8B). Thus, the gain model overestimated the slope of the P1-based CRFs in the focused attention condition (compare blue squares and the black dotted curve in the middle panel of Fig 8B; R^2^ = 0.671). Instead, the noise model provided a significantly better fit compared to the gain model (compare red and black dotted lines in Fig 8B) (R^2^ = 0.874, F(1, 8) = 12.99, p = 0.007, nested test). In addition, the model estimated that the noise parameter (σ) had to be reduced by 31.67%. This reduction of ~32% is roughly analogous to those observed in single-unit and multiple-unit recording data acquired with highly trained NHPs (~50% reduction) (33,38).

**Fig 8.**
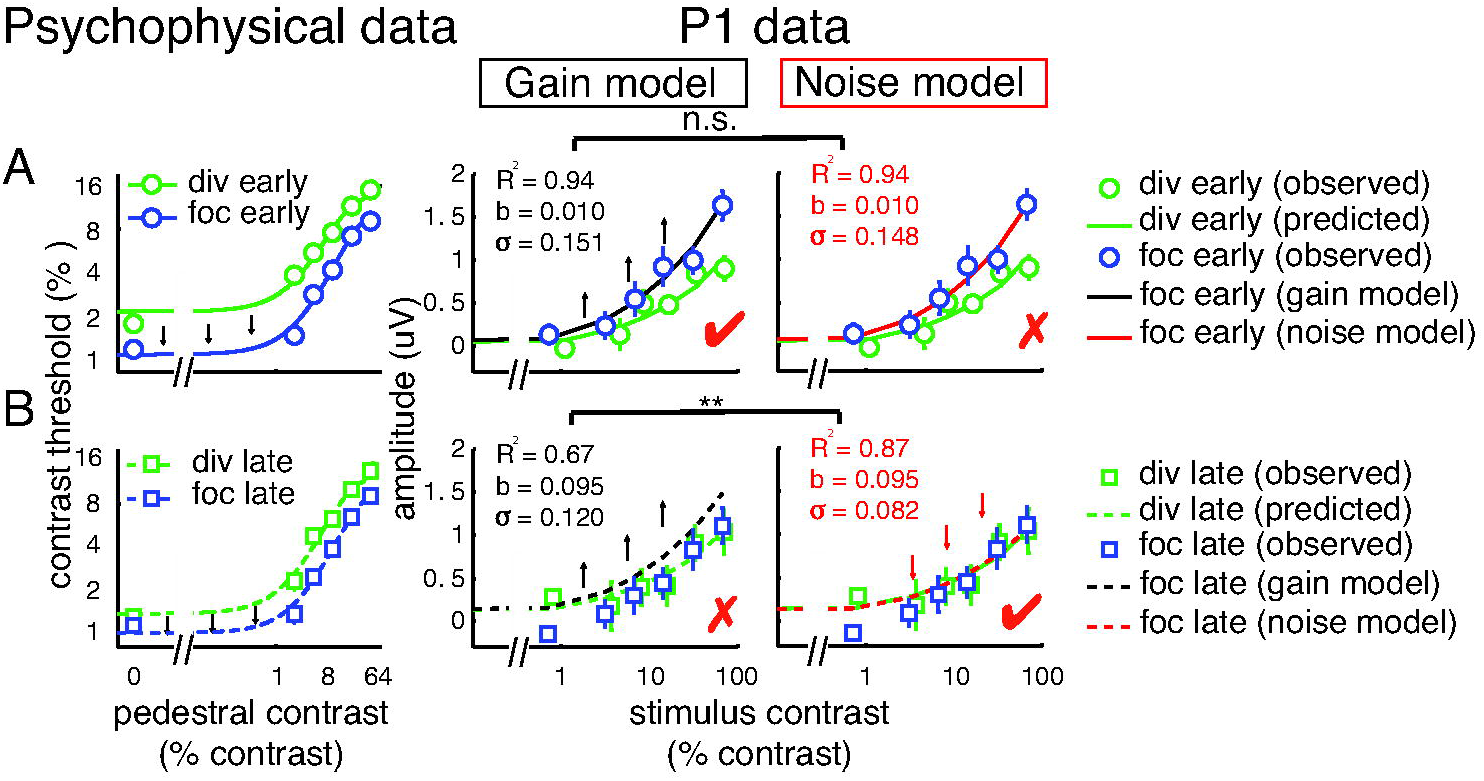
Linking changes in the psychophysical data and the P1 data using gain and noise models. (A) A quantitative model based on the signal detection theory (SDT) reveals that, early in training, attention-induced improvements in behavioral performance is sufficiently explained by the gain model, and the noise model does not significantly improve the fit. (B) In contrast, later in training, the noise model provides a significantly better prediction than the gain model late in training.

In addition to attentional gain and noise reduction, attention can also impact behavior by enhancing the efficiency with which sensory responses are read-out by later sensorimotor and decision-related mechanisms (25,26,59,60). Therefore, we also considered a variant of an efficient read-out model that is based on a max-pooling rule (Equation 11) to account for behavior during the late training phase (25,26). However, given that the noise and gain models almost perfectly predicted behavior, using efficient read-out actually impaired model predictions in this data set (Fig 9). Collectively, these results suggest that training reduced the impact of gain mechanisms and that noise modulations gradually come to play a more dominant role in predicting behavior.

**Fig 9.**
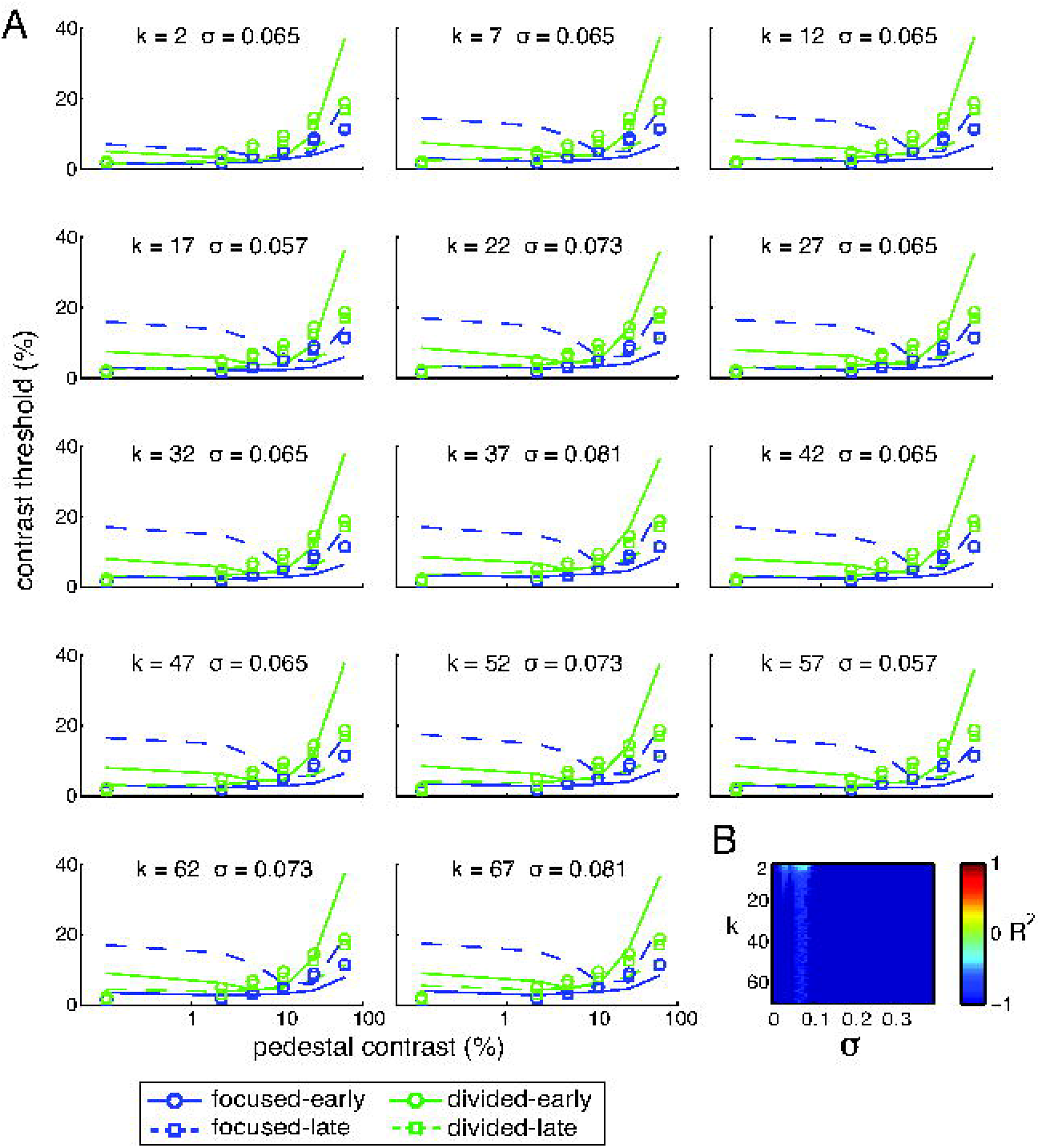
Linking changes in the psychophysical data and the P1 data using the efficient read-out model. (A) The psychophysical contrast-discrimination thresholds estimated from the P1 data using the max-pooling rule (Equation 11) with different *k* values and with the noise values that yield the best fits for those *k* values. (B) Corresponding R^2^ values. Overall, the efficient read-out model did not accurately capture the link between behavior and the P1 data, as R^2^ values are below zero for all *k* and noise values.

### Attentional gain amplification of the posterior late positive deflection (LPD) component persisted throughout training

In addition to our main analysis of attentional modulations of the P1 component, we also examined attentional modulations of another ERP component referred to as the late positive deflection (LPD or P3), which emerged in a later time window about 230-380ms in more central posterior electrodes (Fig 4). This ERP component is thought to index more prolonged post-sensory processes such as sensory evidence accumulation during decision making (14,45,61–64). As shown in Fig 10A, we found that that attentional gain amplification in the LPD occurred to a comparable degree across training phases. For both early and late training phases, we observed significant main effects of attention (p < 0.001 and p = 0.01 for early and late; resampling test, two-tailed) and stimulus type (p <0.001 and p = 0.003 for early and late) and significant interactions between the two factors on the maximum response (p < 0.001 and p =0.035 for early and late; Fig 10B top). These modulations were driven by an increase in the maximum response in the focused target condition relative to all the other conditions (all p’s < 0.001, Holm-Bonferroni correction, two-tailed). Importantly, we observed no significant changes in the maximum response across early and late training phases in any experimental condition (all p’s ≥ 0.21, two-tailed). For the contrast at which the response reached half maximum, we also found significant main effects of attention (p = 0.010) and stimulus type (p = 0.004), as well as a significant interaction between these factors (p = 0.001, resampling test) (Fig 10B bottom). These results were driven by an increase in the contrast at which the response reached half maximum (decrease in contrast gain) in the focused non-target condition compared to all the other conditions (all p’s ≤ 0.002, Holm-Bonferroni correction, two-tailed). Also, we observed no significant changes in the contrast at which the response reached half maximum across early and late training phases in any experimental condition (all p’s ≥ 0.26, two-tailed). Additional analyses also revealed that attentional gain amplification in the LPD occurred to a comparable degree when the data are divided into 4 and 10 training phases (S1 Fig 3).

**Fig 10.**
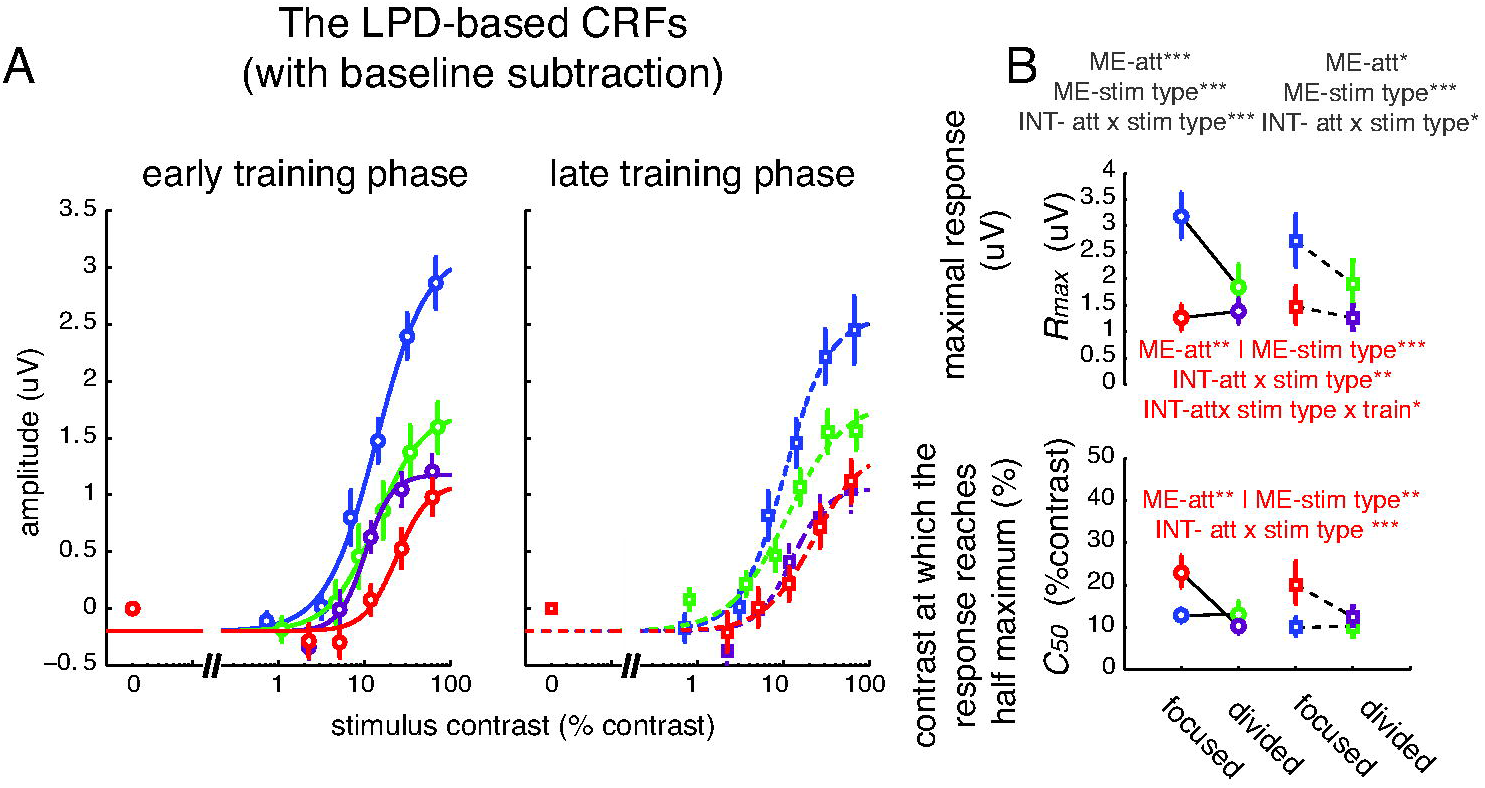
The LPD component with baseline subtraction. (A) The CRF based on the amplitude of the LPD component averaged over posterior electrodes from 230-380ms post-stimulus. Focused attention resulted in increased gain amplification of the LPD during both early and late training phases. (B) Corresponding maximal response and contrast at which response reaches half maximum for the LPD-based CRFs. Error bars in (A) represent within-subject SEM. Error bars in (B) represent the 68% CIs. *, **, and *** present significant main effects (ME) and interactions (INT) with p <0.05, p<0.01 and p <0.001, respectively. Red signs show significance when data were compared across training phases. Gray signs show significance when data were compared within each training phase. See data divided into 4 and 10 training phases in S1 Fig 3 and data without baseline subtraction in Fig 11. Note that the contrast values on the x-axis are not exactly the same across target and non-target conditions because, in the target conditions, we used the averaged contrast values between the pedestal and incremental stimuli.

For comparison, we also analyzed the LPD data without subtracting the baseline activity levels. As illustrated in Fig 11, there were robust modulations of the baseline activity of the LPD component. A nested model comparison analysis confirmed that allowing the baseline parameter to change freely significantly improved the goodness of fit compared to fixing the baseline parameter across experimental conditions (F(7, 15) = 18.03, p <0.001, nested-test). An additional resampling analysis revealed a significant main effect of stimulus type (p < 0.001) and a significant interaction of stimulus type and attention on the baseline parameter (p = 0.013) on the baseline parameter (Fig11B right). This was driven by a significantly larger difference in the baseline activity between the focused non-target and the focused target conditions compared to the difference in baseline activity between the divided target and divided non-target conditions (p = 0.013, two-tailed). We speculate that this elevated baseline response in the focused non-target condition reflects the influence from the decision-related processes associated with the target stimuli, which were enhanced by focused attention. Note that the baseline elevation in the focused non-target condition occurred in an additive fashion because the focused non-target stimuli of each contrast level were simultaneously presented along with focused target stimuli of all possible contrast levels. Therefore, on average, the responses associated with the focused non-target stimuli of each contrast level were contaminated by a similar amount by responses associated with the focused target stimuli averaged across all contrast values, and these effects of stimulus pairing were removed via the baseline subtraction method in the previous section (Fig 10). In addition, we observed a significant interaction between attention and training on the baseline parameter (p = 0.043). This was driven by an increase in the baseline activity in the divided attention conditions in the late training phase compared to the early training phase (p = 0.010, two-tailed), without changes in the focused attention conditions (p = 0.57, two-tailed). This result is consistent with the behavioral result where significant training effects were only observed in the divided attention condition, particularly for low contrast levels, but not in the focused attention condition (Fig 3B).

**Fig 11.**
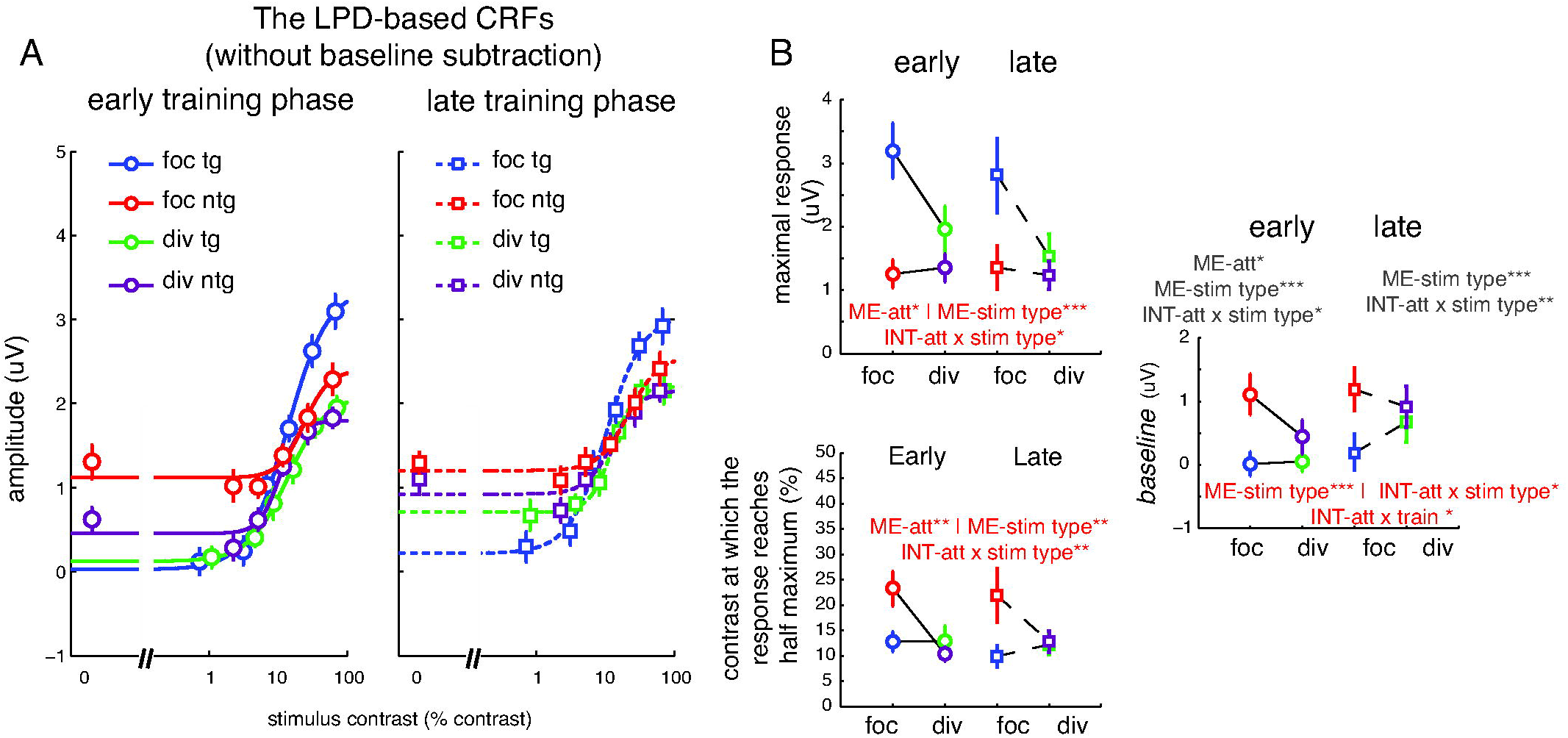
The LPD component without baseline subtraction. (A) The LPD CRFs, based on ERPs without baseline subtraction. (B) Corresponding maximal response, contrast at which response reaches half maximum, and baseline parameters. Error bars in (A) represent within-subject SEM. Error bars in (B) represent the 68% CIs. *, **, and *** present significant main effects (ME) and interactions (INT) with p <0.05, p<0.01 and p <0.001, respectively. Red signs show significance when data were compared across training phases. Gray signs show significance when data were compared within each training phase. Note that the contrast values on the x-axis are not exactly the same across target and non-target conditions because, in the target conditions, we used the averaged contrast values between the pedestal and incremental stimuli.

While the LPD results without baseline subtraction revealed robust modulations in the baseline parameters of the LPD-based CRFs, overall modulations of other parameters including the maximum response and the contrast at which the response reached half maximum (Fig 11B left) are consistent with the results with the subtraction method (Fig 10B). Specifically, we observed robust response and contrast gain modulations across early and late training phases. This is statistically confirmed by significant main effect of attention (p’s = 0.011 and 0.010) and stimulus type (p’s < 0.001 and = 0.008) and a significant interaction between the two factors on the maximum response and the contrast at which the response reached half maximum, respectively (p’s = 0.012 and p = 0.003). Overall, the attention effects on the LPD, which persisted across training phases, stand in contrast to the attention effects on P1 results, which were significantly attenuated with training.

## Discussion

Recent studies using quantitative modeling suggest that there are several candidate mechanisms that can link attentional modulations in visual cortex with attentional modulations of behavior (14,25,26,33,50). While some discrepancies between the putative mechanisms are likely due to differences in stimulus display properties and task designs (15,41,65–67) and methods of measuring neural activity (14,42,43,65), we show here that another major factor is the duration of training. Going beyond previous studies of training that have shown changes in attentional modulations, we quantitatively modeled the relationship between neural activity and behavior to systematically examine the relative contributions of different attention mechanisms (68–72). We found that attentional gain of the visual P1 component accurately predicted attention-induced behavioral benefits early in training. However, this attentional gain was abolished later in training and our SDT-based model (14,25,26,43,48,49) suggested that a substantial reduction in noise was required to explain behavior. In contrast to the P1 results, a similar amount of attentional gain in the LPD component persisted across early and late in training. On a more general level, the observation that the link between attentional modulations of a well-established neural marker like the P1 and behavior can qualitatively change over time underscores the importance of critically considering training when generalizing empirical observations across methods and model systems. This is particularly important given the significant discrepancies in the amount of training that different classes of experimental subjects receive.

The observation of attentional gain of the P1 component during the early training phase is consistent with many prior studies in human participants that reported similar attentional gain effects even when subjects were minimally trained (6– 11,13,14,31,46,47). The attenuation of attentional gain with training, however, is more consistent with observations that noise reduction is the dominant form of attentional modulation in highly trained NHPs (33,38). Interestingly, the SDT model that we used in the current study suggests that the degree of noise reduction needed to compensate for the reduction in P1 gain (~32% reduction) is not far from estimates reported in previous monkey neurophysiology studies (~50% reduction) (33,38). Note that even though this P1 attentional gain modulation occurred very early (~100ms post-stimulus), such modulations likely reflect both feedforward and feedback processes (6,9,73).

Consistent with our study, previous perceptual learning studies have also shown that the P1 component and the earlier C1 component (~50ms post-stimulus) undergo a reduction in amplitude after extended training (74,75 but see 76). A previous neuroimaging study also observed a training-related reduction in activity in primary visual cortex (77). In the present study, training specifically reduced P1 amplitude, however; this training induced change in P1 amplitude occurred only in the focused target condition, with no modulations associated with the focused non-target or the divided attention conditions. Therefore, the training-induced changes in early visual signals that we observe here reflect an attentional modulation and not a more general perceptual learning effect that is related just to repeated exposure to the stimulus set.

In contrast to the P1 results, a comparable amount of gain modulations of the LPD component were observed across training phases, even though attentional gain modulation of the P1 was abolished after training. The LPD has been closely linked to post-sensory decision-related processing and is associated with factors such as response confidence, task difficulty, and decision times (14,45,47,61–64,78,79). Recent EEG studies (63,64) have demonstrated that this LPD component also tracks the accumulation of sensory evidence in a manner that is similar to the average response profile of neurons in the lateral intrapareital area and the frontal eye fields in monkeys during decision making (80–88). Collectively, these findings suggest that while the mechanisms that support attentional modulations of early visual signals shift with training, attentional gain modulations of processing that is closer to the ‘read-out’ stage remain relatively stable over time.

In conclusion, our data demonstrate that attentional gain of the visually evoked P1 component plays a prominent role in enhancing perceptual sensitivity early in training, but noise reduction is required after extensive training. In contrast, attentional gain of the LPD component persists throughout training. This pattern is consistent with an attention-related improvement in the efficiency of the transfer of information, such that earlier stages provide more reliable information to downstream decision-related mechanisms after training. Most importantly, our data show that training can qualitatively alter the relationship between attentional modulations of neural responses and behavior, and this observation carries important implications for understanding attention, as well as for linking observations collected from different model systems that may employ substantially different amounts of training (c.f. 44).

## Materials and Methods

### Subjects

Twenty-three neurologically healthy human observers with normal or corrected-to-normal vision were recruited from the University of California, San Diego (UCSD). All participants provided written informed consent as required by the local Institutional Review Board at UCSD. Data from 10 subjects were discarded in the main analysis due to failure to complete the experimental protocol (20 EEG sessions). One subject only completed a behavioral training session. Among the other 9 subjects, 1, 1, 1, 1, 3, 1, and 2 subjects voluntarily withdrew after the 2^nd^, 6^th^, 8^th^, 10^th^, 12^th^, and 14^th^ EEG sessions. In addition, one of 13 subjects who completed 20 EEG sessions was discarded due to excessive small saccades (>90% of trials). This left 12 subjects in the main analysis (7 female, 20-26 years old, all right-handed). Subjects were compensated at a rate of 10 and 15 US dollars per hour for behavioral training and EEG sessions, respectively.

### Stimuli and task

Stimuli were presented on a PC running Windows XP using MATLAB (Mathworks Inc., Natick, MA) and the Psychophysics Toolbox (version 3.0.8) (89,90). Participants were seated 60 cm from the CRT monitor (which had a grey background of 34.51 cd/m^2^, 60Hz refresh rate) in a sound-attenuated and electromagnetically shielded room (ETS Lindgren).

We employed a 2-IFC contrast discrimination task (Fig 2A). Each trial started with a colored cue (pre-cue) instructing subjects where to attend on each trial. A red cue corresponded to the lower left quadrant, a blue cue corresponded to the lower right quadrant (focused attention conditions), and a green cue indicated that subjects should attend both lower quadrants (divided attention condition). The focused attention cue was 100% valid, whereas the divided attention cue was 50% valid (i.e. the target was equally likely to be presented in the left or the right quadrant). The pre-cue appeared for 500ms followed by a 400-600ms blank inter-stimulus interval (ISI). This ISI was followed by two successive stimulus presentations (the first and second stimulus intervals) with each presentation containing a pair of sinusoidal Gabor stimuli (spatial frequency, 1.04 cycles/°; SD of Gaussian window, 1.90°) located in the lower left and right quadrants (±8.58° and −7.63° from the horizontal and vertical meridians, respectively). Each pair of stimuli appeared for 300ms, followed by a 600-800ms ISI (pseudo-randomly jittered from a uniform distribution). The pedestal contrasts of the Gabor stimuli were randomly selected from six values: 0, 2.24, 5.13, 11.75, 26.92, and 61.66% Michelson contrasts. Contrast values, except for 0%, were jittered ±0.01 log contrast from the mean contrast value. The orientations of the left and right Gabors were identical within each trial, and the orientation value on a given trial was randomly drawn from a uniform distribution. During one of the two stimulus intervals, a contrast increment (Δc) was added to either the left or the right Gabor stimulus for the entire duration of that interval. After the second stimulus interval, the post-cue appeared to inform subjects whether the left (a red cue) or the right stimulus (a blue cue) contained this contrast increment. Subjects reported whether the increment occurred during the first or the second stimulus interval and they were told to prioritize accuracy; there was no response deadline.

On the first day, subjects participated in a ~2.5h behavioral training session where a staircase procedure (3-down, 1-up) was applied to estimate the contrast discrimination thresholds for each attention condition and each pedestal contrast level (see a similar method in 14). These thresholds were then used in the first EEG session. Subjects completed 20 EEG sessions (1-2 sessions each day and 2-3 days a week). Each EEG session contained a total of 8 experimental blocks and contained 288 trials, where all experimental conditions were counterbalanced: 2 (attention conditions: focused, divided) × 2 (target locations: left, right) × 2 (target intervals: first, second) × 6 (pedestal contrast levels of target) × 6 (pedestal contrast levels of non-target). The contrast threshold (Δc) for each attention condition and each target pedestal contrast was adjusted after each EEG session so that accuracy was maintained at ~76% (d’ = ~1) across all experimental conditions. Across the 12 subjects, the average time elapsed between the initial behavioral training session and the 1^st^ EEG session, between the 1^st^ and the 11^th^ EEG sessions (early-training phase), and between the 11^th^ and the last EEG sessions was 3.33 ± 0.66, 17.41 ± 2.31, and 11.50 ± 1.12 days (mean ± SEM), respectively (Fig 2B).

### Behavioral analysis

Contrast discrimination thresholds were measured at ~76% hit rate (d’ = ~1) for all attention conditions (focused and divided attention), training phases (early and late), and stimulus pedestal contrasts (0% to 61.66%). The within-subject SEM of the data for each contrast level was calculated using the Loftus and Masson method (91). Specifically, the mean value between attention and training conditions was removed from the individual subject data before computing the SEM for each contrast value. Three-way repeated measures ANOVAs with within-subject factors of attention condition, training phase, and stimulus pedestal contrast were performed to test the main effect of each of these factors and their interactions on contrast discrimination thresholds. Post-hoc paired t-tests were then used to examine attention effects and learning effects on the contrast discrimination threshold data for each pedestal contrast level (one-tailed, corrected by Holm-Bonferroni method). We used one-tailed statistics here, under the assumption that perceptual sensitivity is enhanced with focused attention and with training.

### EEG preprocessing and analysis

We recorded EEG data with a 64+8 channel Biosemi ActiveTwo system at a 512-Hz sampling rate. All signal offsets from the CMS-DRL reference were maintained below 20 uV. We employed EEGlab11.0.3.1b (92) and custom MATLAB scripts to preprocess the EEG data offline. First, we re-referenced the continuous EEG data to the mean of the two mastoid electrodes and applied 0.25-Hz high-pass and 55-Hz low-pass Butterworth filters (3rd order). Second, the data were segmented into epochs extending from 500ms before to 3500ms after the trial onset. Third, prominent eye blink artifacts were first rejected by independent component analysis (93). We then discarded epochs contaminated by residual eye blinks and vertical eye movements (more than ±80-120 μV deviation from zero, with thresholds chosen for each individual subject), horizontal eye movements (more than ±75-90 μV deviation from zero), excessive muscle activity, or drifts using threshold rejection and visual inspection (11.23 % of trials ± 1.74% S.E.M). Lastly, the data were aligned to the stimulus onset and baseline-corrected based on the mean response from 0-200ms before stimulus onset. For all individual subjects, eye bias scores computed by the difference between averaged horizontal electrooculography contralateral and ipsilateral to the stimulus of interest divided by two are less than 1.6 μV, corresponding to less than 0.1° visual angle which is a standard criterion used in ERP studies (94). Moreover, no difference in eye bias scores were observed across early and late training phases (S1 Fig 4). These results support the notion that any residual horizontal eye movements did not contaminate training-related changes in ERPs.

The artifact-free EEG data were then sorted into the following bins: 2 attention conditions (focused and divided attention) x 2 stimulus types (target and non-target) x 2 training phases (early and late) x 6 stimulus contrast levels x 2 stimulus intervals (first and second) x 2 stimulus locations (left and right). Note that in Supporting Information (S1 Figs 3&4) the data are also sorted into 4 and 10 training phases. The stimulus-locked ERPs were then computed by averaging the EEG data in each bin. To extract ERPs evoked by the stimulus of interest (i.e., subtract out responses evoked by stimuli paired with the stimulus of interest) and minimize confounds from any anticipatory effect from the cue, we subtracted the ERPs evoked by the pedestal 0% contrast stimulus (i.e., when no stimulus was present in the contralateral visual field with respect to a given EEG electrode) from the ERPs on all other conditions (11,14,57) (Fig 4). Thus, the response that was subtracted should be interpreted as “the mean response evoked by an ipsilateral stimulus when no stimulus was presented in the contralateral visual field”, and this served to help isolate the ERP specifically associated with the presentation of a contralateral stimulus. It is critical to isolate the CRFs evoked by the stimuli of interest (focused target, focused non-target, divided target, or divided non-target) from the stimuli on the opposite side that were simultaneously presented on the other side of the display (e.g., if the stimulus of interest was a focused target then the stimulus that was paired with it would be a focused non-target). Thus, subtracting out the small response evoked by the ipsilateral stimulus helps to improve the spatial selectivity of the ERP responses. Moreover, without subtracting this 0%-contrast ERP out, it is possible that the attentional modulations that we may observe would be confounded by cue-related and non-spatially selective anticipatory responses rather than attentional modulations of stimulus-evoked responses (i.e. changes in arousal but not changes in selective spatial attention). While studying such non-selective modulations is potentially interesting in its own right, it would complicate the interpretation of the observed attentional modulations of stimulus-evoked responses. This is a serious issue that has been brought up and dealt with in a similar manner in many previous studies that use both EEG and fMRI (11,14,26,57). By subtracting this 0%-contrast ERP out, we controlled for this potential confound. That said we also included the results without this baseline subtraction for comparisons (Figs 7&11; and see detailed methods in later paragraphs).

The mean amplitude of the visual P1 component from 80-130ms post-stimulus was computed across the contralateral-posterior electrodes, where the P1 mean amplitude averaged across all experimental conditions is maximal (PO7, P5, and P7 for the left hemisphere and PO8, P6, and P8 for the right hemisphere). The selected temporal window was based on previous ERP studies of visual attention (8,10,13,14,46,47,95) and the 50ms window size is suggested as the standard by Luck (94). The mean P1 amplitude was then plotted as a function of stimulus contrast to yield the P1-based CRF separately for each attention condition, each stimulus type, and each training phase. On the y-axis, the stimulus contrast values for the focused and divided non-targets were fixed at 0, 2.24, 5.13, 11.75, 26.92, and 61.66% Michelson contrasts. However, since the target sequence contained both pedestal and increment stimuli, we used the averaged contrast values between the two stimuli for plotting the CRFs in the focused and divided target conditions. The within-subject SEM of the data for each contrast level was calculated using the Loftus and Masson method (91) in which the mean value between attention, stimulus type, and training conditions was removed from individual data before computing the SEM for each contrast value. Next, the P1 data were bootstrapped by resampling subjects, with replacement, 10,000 times. In each bootstrap iteration, the CRF data for each attention condition, stimulus type, and training phase were fit with a Naka-Rushton equation:

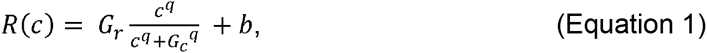

where *R(c)* is the P1 amplitude as a function of stimulus contrast, *G_r_* is a multiplicative response gain factor that controls the vertical shift of the CRF, *G_c_* is a contrast gain factor that controls the horizontal shift of the CRF, *b* is the response baseline offset and *q* is the exponent that controls the speed at which the CRF rises and reaches asymptotes. Given that past EEG studies of spatial attention have consistently reported no changes in response baseline of EEG-based CRFs (3,14–17,29) and in the present study the evoked response to 0%-contrast stimuli was subtracted from all trials, such that the ERP was flat on 0% contrast trials, *b* was fixed as the average of the minimum amplitude across all experimental conditions. We then used a least square error estimation method (fminsearch function in MATLAB) to estimate the maximum response (the response at 100% contrast minus baseline), the contrast at which the response reached half maximum, and the exponent (*q*) parameters. Since in many experimental conditions the CRFs did not saturate at the maximal contrast level (100%), we constrained the fitting procedure so that the maximum response value could not exceed the 1.5 x responses at the 61.66% contrast value (the highest contrast in the stimulus set). *G_r_ and G_c_* were constrained so that they could not be less than 0 and 1, respectively. The exponent *q* was also constrained within a range of −10 to 10. We used the 30% contrast value (about half of 61.66% contrast) as the initial seed value for *G_c_*, the difference between maximum and minimum responses as the seed value for *Gc*, and 1 and 5 for the seed values of the exponent *q* when fitting the CRFs based on the P1 and the LPD (see below for LPD), respectively. The initial seed values for the exponent *q* were adopted from the estimated values based on a previous study (14). To test all main effects and interactions between attention, training and stimulus type, we first computed bootstrap distributions of the differences between the estimated fit parameters:

i. Training effect: early minus late training phases
ii. Attention effect: focused attention minus divided attention
iii. Stimulus type effect: target minus non-target
iv. Two-way interaction between training and attention: (focused attention minus divided attention during the early training phase) minus (focused attention minus divided attention during the late training phase)
v. Two-way interaction between training and stimulus type: (target minus non-target during the early training phase) minus (target minus non-target during the late training phase)
vi. Two-way interaction between attention and stimulus type: (focused target minus focused non-target) minus (divided target minus divided non-target)
vii. Three-way interaction between training, attention, and stimulus type: [(focused target minus focused non-target) minus (divided target minus divided non-target) during the early training stage] minus [(focused target minus focused non-target) minus (divided target minus divided non-target) during the late training stage]

Then, we computed the percentage of values in the tail of each of these compiled distributions that were larger or smaller than zero (two-tailed). If there was any significant interaction between training and any of other remaining factors for each training phase, we then tested for main effects of attention and target type as well as for an interaction between attention and target type using the procedure described above. Post-hoc pairwise comparisons were subsequently performed between these pairwise comparisons and were corrected for multiple comparisons using Holm-Bonferroni method (two-tailed).

For the ‘isolated’ LPD component, the mean amplitude from 230-380ms post-stimulus was computed across the posterior and posterior-occipital electrodes (P5, P7, PO7, P1, Pz, P2, P6, P8, PO8). This analysis window was selected based on the broad activation of the LPD amplitude averaged across all experimental conditions and stimulus contrast levels. Also note that the analysis windows of both P1 and LPD components were chosen to minimize contaminations of the negative-going N1 component that emerges ~150-200ms post-stimulus (see the zoom-in ERP traces for the P1 and LPD components in Figs 5B&5C where minimal negative potentials were observed across these windows). The same bootstrapping, fitting, and statistical analyses described above were also performed on the LPD data.

For comparison, we also analyzed the P1 and the LPD data without baseline subtraction (see results in Figs 7&11). First, we obtained the P1 and LPD components in the same electrodes and the same temporal windows as described above, and plotted the CRFs based on the P1 and LPD mean amplitudes. However, the baseline subtraction described above was not implemented. Next, we fit the Naka-Rushton equation (Equation 1) to characterize the CRFs, but this time we included an additional free parameter to account for baseline differences between conditions. We then performed a nested model comparison to assess the goodness of fit between the model that allowed baseline parameters to change freely (baseline-free model) and the model that fixed the baseline parameter across all experimental conditions (baseline-fixed model) using the following equation:

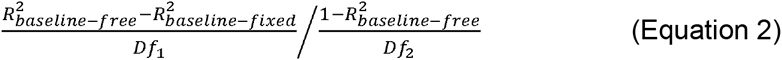

where 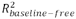 and 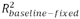 were obtained from the fits of the baseline-free and baseline-fixed models (full and reduced models), respectively. is the number of free parameters in the full model (32: 8 *b’s,* 8 *G_r_’s,* 8 *G_r_’s*, and 8 *q’s* for the focused/divided, target/non-target conditions in the early and late training phases) minus the number of free parameters in the reduced model (25: 8 *G_r_’s,* 8 *G_r_’s*, and 8 *q’s* for the focused/divided, target/non-target conditions in the early and late training phases and 1 *b* shared across all 8 experimental conditions). is the number of observations (48: 6 contrast levels times 8 experimental conditions) minus the number of free parameters in the full model (32) minus 1. The *F* distribution was used to estimate the probability that the full model differed significantly from the reduced model. For the P1 data, the baseline-free model was not significantly better than the baseline-fixed model (see Results), so we only evaluated the significance of the best fit parameters estimated using Equation 1 with a fixed baseline parameter. On the other hand, the baseline-free model was significantly better for the LPD data, so we reported statistical results using a version of Equation 1 with a freely optimized baseline parameter.

### Modeling methods

We adopted a previously established model based on SDT (14,26,42,48) to determine the degree to which attentional gain and noise reduction are needed to explain the relationship between attentional modulations in the psychophysical and ERP data during the early and late training phases. This modeling framework is based on the assumption that perceptual sensitivity (*d’*) is limited by the differential mean response: *R(c+Δc(c))-R(c)* or ΔR, evoked by two different stimuli (i.e. standard and test stimuli) divided by the trial-by-trial variability of those responses (σ):

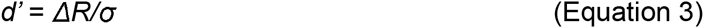

where *R* is the hypothetical CRF estimated using the Naka-Rushton equation (Equation 1). With the combination of the d’ and Naka-Rushton equations, the contrast discrimination thresholds could be estimated based on the derivative (or slope) of the CRF as expressed in the following equation:

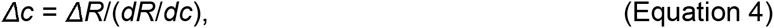

where *dR/dc* is the derivative of the underlying CRF (48).

According to the attentional gain model, attention-induced reductions in contrast discrimination thresholds can be fully explained by an increase in the slope of the ERP-based CRF (*dR/dc*), under the assumption that the neuronal noise (σ) is constant (gain model). In the case where the amount of increase in the CRF slope is insufficient to explain shifts in the psychophysical TvC functions, the σ parameter must be reduced to explain changes in psychophysical contrast thresholds (noise model).

We applied this model using the following procedure: we first estimated the psychophysical TvC functions for the divided attention and focused attention conditions for both the early and late training phases using a polynomial function with least square error estimation methods (fminsearch function in MATLAB). For each training phase, we used the combination of the Naka-Rushton and d’ equations to simulate the CRFs based on the P1 amplitude in the divided target condition. Specifically, the fitting routine started by setting the first point of the estimated CRF (*c_0_* = 0%) to be a baseline parameter (*b*) as the following:

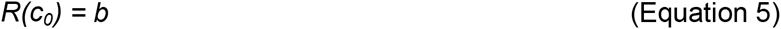

The next contrast (c_1_) was then defined as:

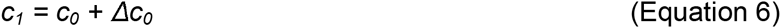

where Δ*c_0_* is the contrast threshold at 0% contrast. Accordingly, the response at *c_1_* was estimated using the d’ equation (Eq2) as:

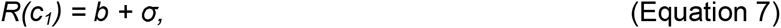

given that *d’* = 1. The next contrast was defined the same way as the following:

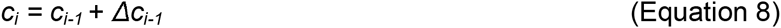

where *i* is the current iteration that is > 1. The response at *c_i_* is then estimated as

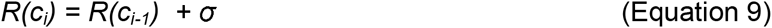

These last two steps (Equations 8&9) were continued until the full CRF was estimated. The baseline and noise parameters (*b* and σ) were optimized by minimizing the least-squares errors between the observed and the predicted CRFs based on the P1 amplitude in the divided target condition. To test if attentional gain changes in the P1 CRFs in each training phase could account for changes in the TvC functions, we estimated the P1 CRFs in the focused target condition using the modeling routine described above but with the *b* and σ parameters fixed based on the values obtained from the divided target condition. Next, we tested if allowing changes in the noise parameter across focused target and divided target conditions could significantly improve the prediction of the model based on SDT. To achieve this, we estimated the P1 CRFs in the focused target condition as described above, except that we allowed the σ parameter to vary freely to find the best fit. The R^2^ value obtained from the gain model with the σ parameter fixed across the divided target and focused target conditions (reduced model) was then compared with the R^2^ value obtained using the noise reduction model in which the σ parameter was also allowed to vary freely across attention conditions (full model). This comparison was done using a nested F test:

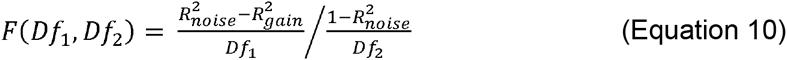

where 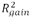 and 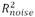 were obtained from the fits of the attentional gain and noise modulation models (reduced and full models), respectively. *Df*_1_ is the number of free parameters in the full model (3: σ for focused attention, σ for divided attention, and *b* shared across attention conditions) minus the number of free parameters in the reduced model (2: σ and *b* shared across attention conditions). is the number of observations (12: 6 contrast levels times two attention conditions) minus the number of free parameters in the full model (3) minus 1. The *F* distribution was used to estimate the probability that the full model differed significantly from the reduced model.

In addition to the gain and noise models based on SDT, we also adopted a variant of an efficient read-out model to see how well it could explain the link between attentional modulations in the P1 component and behavioral data across training stages. To start the procedure, we first fit the neural CRFs based on the P1 amplitudes with the Naka-Rushton equation (Equation 1 and see the fitting procedure below the equation). Since the model requires all responses to be positive values (due to the *k* exponent in the max pooling rule see Equation 11 below), we subtracted the baseline values from the interpolated CRFs of all attention conditions and training stages. Next, for each attention condition of each training phase, we simulated the performance of an ideal observer in 72,000 randomly generated trials, which consisted of 12,000 trials of each of the 6 levels of target pedestal contrasts. These 12,000 trials included 2,000 trials of each of the 6 levels of non-target contrasts. For each simulated trial, we determined the response of each stimulus type (target or non-target) and stimulus interval (the interval that contains the test contrast or pedestal contrast) as a random draw from a Gaussian distribution with mean values equal to the mean amplitude of the interpolated P1 CRFs at the corresponding contrast value. The standard deviation (SD) of the Gaussian distribution is the noise parameter in the d’ equation (Equation 3) and it was varied from 0.001 to 0.393 in 50 0.008-unit incremental steps. Next, the target and non-target related responses (*R_tg_* and *R_ntg_*) were pooled into a single response (*R_p_*) using the max-pooling equation (14,25,26).

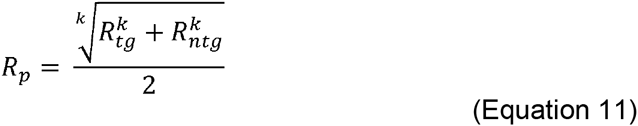

where *k* is an exponent that weights responses to each stimulus in a given interval. Under the assumption that an ideal observer would select the interval that contained a larger pooled response as the interval that contained the incremental target stimulus, we searched for the contrast increment value that yielded 76% accuracy rate across the 12,000 simulated trials at each pedestal contrast level. Here, *k* was varied from 2 to 70 in 69 1-unit incremental steps.

## Acknowledgement

We thank Annalisa Salazar, Sean Deering, Cecillia Chow, and Jhankhana Jani for help with data collection, and Edward Awh, Thomas Sprague, Vy Vo, and Rosanne Rademaker for useful discussions. Supported by NIH R01-092345 to J.T.S, a James S. McDonnell Foundation award to J.T.S, and by an HHMI International Fellowship to S.I.

## References

1. Buracas GT, Boynton GM. The Effect of Spatial Attention on Contrast Response Functions in Human Visual Cortex. J Neurosci [Internet]. 2007;27(1):93–7. Available from: http://www.jneurosci.org/cgi/doi/10.1523/JNEUROSCI.3162-06.2007

2. Connor CE, Preddie DC, Gallant JL, Van Essen DC. Spatial attention effects in macaque area V4. J Neurosci. 1997;17(9):3201–14.

3. Di Russo F, Spinelli D, Morrone MC. Automatic gain control contrast mechanisms are modulated by attention in humans: Evidence from visual evoked potentials. Vision Res. 2001;41(19):2435–47.

4. Gouws AD, Alvarez I, Watson DM, Uesaki M, Rogers J, Morland AB. On the Role of Suppression in Spatial Attention: Evidence from Negative BOLD in Human Subcortical and Cortical Structures. J Neurosci [Internet]. 2014;34(31):10347–60. Available from: http://www.jneurosci.org/cgi/doi/10.1523/JNEUROSCI.0164-14.2014

5. Haenny PE, Maunsell JH, Schiller PH. State dependent activity in monkey visual cortex. II. Retinal and extraretinal factors in V4. Exp brain Res. 1988;69(2):245– 59.

6. Noesselt T, Hillyard SA, Woldorff MG, Schoenfeld A, Hagner T, Jäncke L, et al. Delayed striate cortical activation during spatial attention. Neuron [Internet]. 2002;35(3):575–87. Available from: http://www.sciencedirect.com/science/article/B6WSS-46FS7D0-K/2/164c7b333f5a4ed58c6dbe3c8e6f3098

7. Hillyard S a, Vogel EK, Luck SJ. Sensory gain control (amplification) as a mechanism of selective attention: electrophysiological and neuroimaging evidence. Philos Trans R Soc Lond B Biol Sci. 1998;353(1373):1257–70.

8. Mangun GR, Hillyard S a. Allocation of visual attention to spatial locations: tradeoff functions for event-related brain potentials and detection performance. Percept Psychophys. 1990;47(6):532–50.

9. Hillyard S a, Anllo-Vento L. Event-related brain potentials in the study of visual selective attention. Proc Natl Acad Sci U S A. 1998;95(3):781–7.

10. Mangun GR, Hillyard SA. The spatial allocation of visual attention as indexed by event-related brain potentials. HumFactors. 1987;29(2):195–211.

11. Störmer VS, McDonald JJ, Hillyard S a. Cross-modal cueing of attention alters appearance and early cortical processing of visual stimuli. Proc Natl Acad Sci U S A. 2009;106(52):22456–61.

12. Voorhis S, Hillyard S a. Visual evoked potentials and selective attention to points in space. Percept Psychophys [Internet]. 1977;22(1):54–62. Available from: http://www.springerlink.com/index/10.3758/BF03206080

13. Mangun GR, Hillyard S a. Spatial gradients of visual attention: behavioral and electrophysiological evidence. Electroencephalogr Clin Neurophysiol. 1988;70:417–28.

14. Itthipuripat S, Ester EF, Deering S, Serences JT. Sensory gain outperforms efficient readout mechanisms in predicting attention-related improvements in behavior. J Neurosci [Internet]. 2014;34(40):13384–98. Available from: http://www.ncbi.nlm.nih.gov/pubmed/25274817

15. Itthipuripat S, Garcia JO, Rungratsameetaweemana N, Sprague TC, Serences JT. Changing the spatial scope of attention alters patterns of neural gain in human cortex. J Neurosci [Internet]. 2014;34(1):112–23. Available from: http://www.ncbi.nlm.nih.gov/pubmed/24381272

16. Kim YJ, Grabowecky M, Paller K a, Muthu K, Suzuki S. Attention induces synchronization-based response gain in steady-state visual evoked potentials. Nat Neurosci. 2007;10(1):117–25.

17. Lauritzen TZ, Ales JM, Wade AR. The effects of visuospatial attention measured across visual cortex using source-imaged, steady-state EEG. J Vis. 2010;10(14):1–17.

18. Lee J, Maunsell JHR. A normalization model of attentional modulation of single unit responses. PLoS One. 2009;4(2):e4651.

19. Lee J, Maunsell JHR. The effect of attention on neuronal responses to high and low contrast stimuli. J Neurophysiol. 2010;104(2):960–71.

20. Martínez-Trujillo JC, Treue S. Attentional modulation strength in cortical area MT depends on stimulus contrast. Neuron. 2002;35(2):365–70.

21. McAdams CJ, Maunsell JH. Effects of attention on orientation-tuning functions of single neurons in macaque cortical area V4. J Neurosci [Internet]. 1999;19(1):431–41. Available from: http://www.ncbi.nlm.nih.gov/pubmed/9870971

22. Moran J, Desimone R. Selective attention gates visual processing in the extrastriate cortex. Science. 1985;229(4715):782–4.

23. Motter BC. Focal attention produces spatially selective processing in visual cortical areas V1, V2, and V4 in the presence of competing stimuli. J Neurophysiol. 1993;70(3):909–19.

24. Murray SO. The effects of spatial attention in early human visual cortex are stimulus independent. J Vis. 2008;8(10):2.1–11.

25. Hara Y, Gardner JL. Encoding of graded changes in spatial specificity of prior cues in human visual cortex. J Neurophysiol [Internet]. 2014;112(11):2834–49. Available from: http://jn.physiology.org/cgi/doi/10.1152/jn.00729.2013

26. Pestilli F, Carrasco M, Heeger DJ, Gardner JL. Attentional Enhancement via Selection and Pooling of Early Sensory Responses in Human Visual Cortex. Neuron [Internet]. 2011;72(5):832–46. Available from: http://linkinghub.elsevier.com/retrieve/pii/S0896627311008762

27. Reynolds JH, Pasternak T, Desimone R. Attention Increases Sensitivity of V4 Neurons. Neuron [Internet]. 2000;26(3):703–14. Available from: http://www.sciencedirect.com/science/article/pii/S0896627300812064

28. Sprague TC, Serences JT. Attention modulates spatial priority maps in the human occipital, parietal and frontal cortices. Nat Neurosci [Internet]. Nature Publishing Group; 2013;16(12):1–14. Available from: http://dx.doi.org/10.1038/nn.3574%5Cnpapers3://publication/doi/10.1038/nn.3574

29. Wang J, Wade AR. Differential attentional modulation of cortical responses to S-cone and luminance stimuli. J Vis. 2011;11(6):1.

30. Williford T, Maunsell JHR. Effects of Spatial Attention on Contrast Response Functions in Macaque Area V4. 2006;40–54.

31. Woldorff MG, Fox PT, Matzke M, Lancaster JL, Veeraswamy S, Zamarripa F, et al. Retinotopic organization of early visual spatial attention effects as revealed by PET and ERPs. Hum Brain Mapp. 1997;5:280–6.

32. Ruff DA, Cohen MR. Attention can either increase or decrease spike count correlations in visual cortex. Nat Neurosci [Internet]. 2014;17(11):1591–7. Available from: http://www.nature.com/doifinder/10.1038/nn.3835

33. Cohen MR, Maunsell JHR. Attention improves performance primarily by reducing interneuronal correlations. Nat Neurosci [Internet]. 2009;12(12):1594–600. Available from: http://www.nature.com/doifinder/10.1038/nn.2439

34. Cohen MR, Kohn A. Measuring and interpreting neuronal correlations. Nat Neurosci [Internet]. 2011;14(7):811–9. Available from: http://www.nature.com/doifinder/10.1038/nn.2842

35. Luo TZ, Maunsell JHR. Neuronal Modulations in Visual Cortex Are Associated with Only One of Multiple Components of Attention. Neuron [Internet]. Elsevier Inc.; 2015;86(5):1182–8. Available from: http://www.sciencedirect.com/science/article/pii/S0896627315004146

36. Niebergall R, Khayat PS, Treue S, Martinez-Trujillo JC. Expansion of MT neurons excitatory receptive fields during covert attentive tracking. J Neurosci [Internet]. 2011;31(43):15499–510. Available from: http://www.ncbi.nlm.nih.gov/pubmed/22031896

37. Mitchell JF, Sundberg KA, Reynolds JH. Differential Attention-Dependent Response Modulation across Cell Classes in Macaque Visual Area V4. Neuron [Internet]. 2007;55(1):131–41. Available from: http://linkinghub.elsevier.com/retrieve/pii/S0896627307004497

38. Mitchell JF, Sundberg K a., Reynolds JH. Spatial Attention Decorrelates Intrinsic Activity Fluctuations in Macaque Area V4. Neuron [Internet]. Elsevier Ltd; 2009;63(6):879–88. Available from: http://dx.doi.org/10.1016/j.neuron.2009.09.013

39. Noudoost B, Moore T. Control of visual cortical signals by prefrontal dopamine. Nature [Internet]. Nature Publishing Group; 2011;474(7351):372–5. Available from: http://dx.doi.org/10.1038/nature09995

40. Tremblay S, Pieper F, Sachs A, Martinez-Trujillo J. Attentional Filtering of Visual Information by Neuronal Ensembles in the Primate Lateral Prefrontal Cortex. Neuron [Internet]. 2015;85(1):202–15. Available from: http://linkinghub.elsevier.com/retrieve/pii/S0896627314010733

41. Reynolds JH, Heeger DJ. The Normalization Model of Attention. Neuron [Internet]. Elsevier Inc.; 2009;61(2):168–85. Available from: http://dx.doi.org/10.1016/j.neuron.2009.01.002

42. Hara Y, Pestilli F, Gardner JL. Differing effects of attention in single-units and populations are well predicted by heterogeneous tuning and the normalization model of attention. Front Comput Neurosci [Internet]. 2014;8(February):12. Available from: http://www.pubmedcentral.nih.gov/articlerender.fcgi?artid=3928538&tool=pmcentrez&rendertype=abstract

43. Itthipuripat S, Serences JT. Integrating Levels of Analysis in Systems and Cognitive Neurosciences: Selective Attention as a Case Study. Neurosci [Internet]. 2015; Available from: http://nro.sagepub.com/cgi/doi/10.1177/1073858415603312

44. Birman D, Gardner JL. Parietal and prefrontal: categorical differences? Nat Publ Gr [Internet]. Nature Publishing Group; 2016;19(1):5–7. Available from: http://dx.doi.org/10.1038/nn.4204

45. Hillyard S, Squires K, Baue J, Lindsay P. Evoked potential correlates of response criterion in auditory signal detection. [Internet]. Science (New York, N.Y.). 1972. p. 1357–60. Available from: http://www.ncbi.nlm.nih.gov/pubmed/5035489

46. Johannes S, Münte TF, Heinze HJ, Mangun GR. Luminance and spatial attention effects on early visual processing. Cogn Brain Res [Internet]. 1995;2(3):189–205. Available from: http://www.ncbi.nlm.nih.gov/pubmed/7580401

47. Mangun GR, Buck L a. Sustained visual spatial attention produces costs and benefits in response time and evoked neural activity. Neuropsychologia. 1998;36(3):189–200.

48. Boynton GM, Demb JB, Glover GH, Heeger DJ. Neuronal basis of contrast discrimination. Vision Res. 1999;39(2):257–69.

49. Serences JT. Mechanisms of Selective Attention: Response Enhancement, Noise Reduction, and Efficient Pooling of Sensory Responses. Neuron [Internet]. Elsevier Inc.; 2011;72(5):685–7. Available from: http://dx.doi.org/10.1016/j.neuron.2011.11.005

50. Baruni JK, Lau B, Salzman CD. Reward expectation differentially modulates attentional behavior and activity in visual area V4. Nat Neurosci [Internet]. Nature Publishing Group; 2015;18(11):1656–63. Available from: http://www.nature.com/doifinder/10.1038/nn.4141

51. Gorea a, Sagi D. Disentangling signal from noise in visual contrast discrimination. Nat Neurosci. 2001;4(11):1146–50.

52. Huang L, Dobkins KR. Attentional effects on contrast discrimination in humans: Evidence for both contrast gain and response gain. Vision Res. 2005;45(9):1201– 12.

53. Legge GE, Foley JM. Contrast masking in human vision. J Opt Soc Am [Internet]. 1980;70(12):1458–71. Available from: http://www.ncbi.nlm.nih.gov/pubmed/7463185

54. Ross J, Speed HD, Morgan MJ. The effects of adaptation and masking on incremental thresholds for contrast. Vision Res [Internet]. 1993;33(15):2051–6. Available from: http://www.ncbi.nlm.nih.gov/pubmed/8266646

55. Adini Y, Sagi D, Tsodyks M. Context-enabled learning in the human visual system. 2002;415(February):2–5.

56. Yu C, Klein S a, Levi DM. Perceptual learning in contrast discrimination and the (minimal) role of context. J Vis. 2004;4(3):169–82.

57. Talsma D, Woldorff MG. Selective attention and multisensory integration: multiple phases of effects on the evoked brain activity. J Cogn Neurosci [Internet]. 2005;17(7):1098–114. Available from: http://www.ncbi.nlm.nih.gov/pubmed/16102239

58. Herrero JL, Gieselmann MA, Sanayei M, Thiele A. Attention-Induced Variance and Noise Correlation Reduction in Macaque V1 Is Mediated by NMDA Receptors. Neuron [Internet]. Elsevier Inc.; 2013;78(4):729–39. Available from: http://linkinghub.elsevier.com/retrieve/pii/S0896627313002766

59. Eckstein MP, Peterson MF, Pham BT, Droll J a. Statistical decision theory to relate neurons to behavior in the study of covert visual attention. Vision Res [Internet]. Elsevier Ltd; 2009;49(10):1097–128. Available from: http://www.ncbi.nlm.nih.gov/pubmed/19138699

60. Palmer J, Verghese P, Pavel M. The psychophysics of visual search. Vision Res. 2000;40:1227–68.

61. Squires KC, Hillyard SA, Lindsay PH. Cortical potentials evoked by confirming and disconfirming feedback following an auditory discrimination. Percept Psychophys [Internet]. 1973;13:25–31. Available from: http://www.springerlink.com/index/10.3758/BF03207230

62. Squires NK, Squires KC, Hillyard SA. Two varieties of long-latency positive waves evoked by unpredictable auditory stimuli in man. Electroencephalogr Clin Neurophysiol [Internet]. 1975;38(4):387–401. Available from: http://www.sciencedirect.com/science/article/pii/0013469475902631

63. Kelly SP, O’Connell RG. Internal and External Influences on the Rate of Sensory Evidence Accumulation in the Human Brain. J Neurosci [Internet]. 2013;33(50):19434–41. Available from: http://www.jneurosci.org/cgi/doi/10.1523/JNEUROSCI.3355-13.2013

64. O’Connell RG, Dockree PM, Kelly SP. A supramodal accumulation-to-bound signal that determines perceptual decisions in humans. Nat Neurosci [Internet]. 2012;15(12):1729–35. Available from: http://www.nature.com/doifinder/10.1038/nn.3248

65. Gardner JL. A case for human systems neuroscience. Neuroscience [Internet]. IBRO; 2014;296:130–7. Available from: http://www.ncbi.nlm.nih.gov/pubmed/24997268

66. Herrmann K, Montaser-Kouhsari L, Carrasco M, Heeger DJ. When size matters: attention affects performance by contrast or response gain. Nat Neurosci [Internet]. Nature Publishing Group; 2010;13(12):1554–9. Available from: http://www.nature.com/doifinder/10.1038/nn.2669

67. Zhang X, Japee S, Safiullah Z, Mlynaryk N, Ungerleider LG. A Normalization Framework for Emotional Attention. 2016;1–25.

68. Byers A, Serences JT. Enhanced attentional gain as a mechanism for generalized perceptual learning in human visual cortex. J Neurophysiol [Internet]. 2014;(June):1217–27. Available from: http://www.ncbi.nlm.nih.gov/pubmed/24920023

69. Clark K, Appelbaum LG, van den Berg B, Mitroff SR, Woldorff MG. Improvement in Visual Search with Practice: Mapping Learning-Related Changes in Neurocognitive Stages of Processing. J Neurosci [Internet]. 2015;35(13):5351–9. Available from: http://www.jneurosci.org/cgi/doi/10.1523/JNEUROSCI.1152-14.2015

70. Crist RE, Li W, Gilbert CD. Learning to see: Experience and attention in primary visual cortex. Nat Neurosci [Internet]. 2001;4(5):519–25. Available from: http://www.ncbi.nlm.nih.gov/pubmed/11319561

71. Jehee JFM, Ling S, Swisher JD, van Bergen RS, Tong F. Perceptual Learning Selectively Refines Orientation Representations in Early Visual Cortex. J Neurosci [Internet]. 2012;32(47):16747–53. Available from: http://www.jneurosci.org/cgi/doi/10.1523/JNEUROSCI.6112-11.2012

72. Kelley T a, Yantis S. Neural correlates of learning to attend. Front Hum Neurosci [Internet]. 2010;4(November):216. Available from: http://www.pubmedcentral.nih.gov/articlerender.fcgi?artid=2991198&tool=pmcentrez&rendertype=abstract

73. Lopes da Silva F. EEG and MEG: Relevance to neuroscience. Neuron [Internet]. Elsevier Inc.; 2013;80(5):1112–28. Available from: http://dx.doi.org/10.1016/j.neuron.2013.10.017

74. Pourtois G, Rauss KS, Vuilleumier P, Schwartz S. Effects of perceptual learning on primary visual cortex activity in humans. Vision Res. 2008;48(1):55–62.

75. Zhang G, Cong L-J, Song Y, Yu C. ERP P1-N1 changes associated with Vernier perceptual learning and its location specificity and transfer. J Vis. 2013;13(4):1– 13.

76. Bao M, Yang L, Rios C, He B, Engel SA. Perceptual Learning Increases the Strength of the Earliest Signals in Visual Cortex. J Neurosci [Internet]. 2010;30(45):15080–4. Available from: http://www.jneurosci.org/cgi/doi/10.1523/JNEUROSCI.5703-09.2010

77. Yotsumoto Y, Watanabe T, Sasaki Y. Different Dynamics of Performance and Brain Activation in the Time Course of Perceptual Learning. Neuron. 2008;57(6):827–33.

78. Itthipuripat S, Cha K, Rangsipat N, Serences JT. Value-based attentional capture influences context-dependent decision-making. J Neurophysiol [Internet]. 2015;114(1):560–9. Available from: http://jn.physiology.org/lookup/doi/10.1152/jn.00343.2015

79. Wiener M, Thompson JC. Repetition enhancement and memory effects for duration. Neuroimage [Internet]. Elsevier Inc.; 2015;113:268–78. Available from: http://dx.doi.org/10.1016/j.neuroimage.2015.03.054

80. Ding L, Gold JI. Neural correlates of perceptual decision making before, during, and after decision commitment in monkey frontal eye field. Cereb Cortex. 2012;22(5):1052–67.

81. Gold JI, Shadlen MN. The neural basis of decision making. Annu Rev Neurosci [Internet]. 2007;(30):535–61. Available from: http://www.annualreviews.org/doi/pdf/10.1146/annurev.neuro.29.051605.113038%5Cnpapers2://publication/uuid/BD039A59-7F74-44AB-885F-FFAC88E01CF1

82. Huk AC. Neural Activity in Macaque Parietal Cortex Reflects Temporal Integration of Visual Motion Signals during Perceptual Decision Making. J Neurosci [Internet]. 2005;25(45):10420–36. Available from: http://www.jneurosci.org/cgi/doi/10.1523/JNEUROSCI.4684-04.2005

83. Kim JN, Shadlen MN. Neural correlates of a decision in the dorsolateral prefrontal cortex of the macaque. Nat Neurosci. 1999;2(2):176–85.

84. Mazurek ME. A Role for Neural Integrators in Perceptual Decision Making. Cereb Cortex [Internet]. 2003;13(11):1257–69. Available from: http://www.cercor.oupjournals.org/cgi/doi/10.1093/cercor/bhg097

85. Roitman JD, Shadlen MN. Response of neurons in the lateral intraparietal area during a combined visual discrimination reaction time task. J Neurosci. 2002;22(21):9475–89.

86. Shadlen MN, Newsome WT. Neural basis of a perceptual decision in the parietal cortex (area LIP) of the rhesus monkey. J Neurophysiol [Internet]. 2001;86(4):1916–36. Available from: http://www.ncbi.nlm.nih.gov/pubmed/11600651

87. Woodman GF, Kang M-S, Thompson K, Schall JD. The effect of visual search efficiency on response preparation: neurophysiological evidence for discrete flow. Psychol Sci a J Am Psychol Soc / APS [Internet]. 2008;19(2):128–36. Available from: http://www.ncbi.nlm.nih.gov/pubmed/18271860

88. Law C-T, Gold JI. Neural correlates of perceptual learning in a sensory-motor, but not a sensory, cortical area. Nat Neurosci [Internet]. 2008;11(4):505–13. Available from: http://www.nature.com/doifinder/10.1038/nn2070

89. Brainard DH. The Psychophysics Toolbox. Spat Vis [Internet]. 1997;10(4):433–6. Available from: http://bbs.bioguider.com/images/upfile/2006-4/200641014348.pdf

90. Pelli D. The VideoToolbox software for visual psychophysic:transforming numbers into movies. Spat Vis. 1997;10(4):437–42.

91. Loftus GR, Masson, Micael EJ. Using confidence intervals in within-subject designs. 1994;1:476–90. Available from: papers3://publication/uuid/0DFB63AE-D78E-44E2-B870-1341DCA37BBC

92. Delorme A, Makeig S. EEGLAB: An open source toolbox for analysis of single-trial EEG dynamics including independent component analysis. J Neurosci Methods. 2004;134(1):9–21.

93. Makeig S J., Bell A., Jung T-P, Sejnowski TJ. Independent Component Analysis of Electroencephalographic Data. Adv Neural Inf Process Syst. 1996;8:145–51.

94. Luck SJ. An Introduction to the Event-Related Potential Technique. Cambridge, MA: MIT Press; 2005.

95. Mangun GR, Buonocore MH, Girelli M, Jha AP. ERP and fMRI measures of visual spatial selective attention. Hum Brain Mapp. 1998;6(5–6):383–9.

